# 3’UTR-dependent dynamic changes in *TP53* mRNA localization regulate p53 tumor suppressor activity

**DOI:** 10.1101/2022.04.04.487038

**Authors:** Linshan Hu, Sweta Misra, Baktiar Karim, Skyler Kuhn, Jacqueline Salotti, Srikanta Basu, Nancy Martin, Karen Saylor, Peter F. Johnson

## Abstract

The tumor suppressor p53 triggers senescence in response to oncogenic stress in primary cells. However, the mechanisms by which tumor cells retaining p53 bypass senescence are not fully understood. Here we report that p53 cytostatic activity is inhibited in tumor cells by the 3’ untranslated region (3’UTR) of its mRNA, without altering p53 levels. 3’UTR inhibition requires a long U-rich element (URE) and its binding protein, HuR. The 3’UTR excluded *TP53* mRNAs from a perinuclear compartment containing the CK2 kinase, suppressing p53 phosphorylation on an activating CK2 site, Ser392. In primary cells undergoing oncogene-induced senescence and tumor cells treated with genotoxic agents, *TP53* mRNAs became concentrated in the perinuclear cytoplasm, coinciding with p53 phosphorylation and activation by CK2. In both cases, perinuclear re-localization of *TP53* transcripts required AMPK*α*2-dependent HuR nuclear translocation. ATM kinase activity was essential for DNA damage-induced spatial reprogramming of *TP53* mRNAs, likely through phosphorylation and inactivation of MDM2. MDM2 was required for peripheral localization of *TP53* transcripts and negatively regulated levels of the AMPK*α*2 activating kinase, CaMKK*β*. Our findings reveal a critical role for 3’UTR sequences in suppressing p53 protein activity and provide a new mechanistic framework for p53 activation by DNA damaging agents.

## Introduction

Cellular senescence is induced by various stresses such as DNA damage, oxidative stress and oncogenes and acts as a first line defense against cancer^1^. Oncogene-induced senescence (OIS) is implemented in normal cells by dysregulated signaling from mutant RAS and other oncogenes, thus activating the tumor suppressors p53 and RB. Oncogenic stress increases expression of the ARF tumor suppressor, which stabilizes p53 by neutralizing MDM2^2^, a p53 E3 ubiquitin ligase and oncogene. DNA damage elicits reversible proliferative arrest, senescence, or cell death, which are generally dependent on p53. Severe DNA damage induces apoptosis in tumor cells, including those lacking ARF, through DNA damage response (DDR) pathways that are partially distinct from OIS mechanisms.

As a potent activator of growth arrest, senescence, and apoptosis, p53 is tightly regulated at multiple levels^3,4^. Among these is MDM2-dependent proteosomal degradation, which keeps p53 levels low in unstressed cells^5^. Other evidence suggests that the E3 ligase function of MDM2 is not required to suppress physiological p53 activity^6^, and additional regulation is conferred by MDM2 binding to p53 TAD regions to inhibit transactivation^3^. The importance of MDM2-mediated suppression of p53 is highlighted by genetic studies showing that mice lacking MDM2 exhibit embryonic lethality that can be abrogated in part by loss of p53^7,8^. It has been suggested that p53 activity is also constrained by autoinhibition through a C-terminal domain (CTD), although this region has been implicated in both negative and positive regulation of DNA binding^3^ and in controlling binding site specificity^9^. Moreover, p53 binds to chromatin even in unstressed cells^10,11^. p53 is activated at various levels by at least 40 post-translational modifications (PTMs)^3^. These include phosphorylation on several sites in the TAD regions by the DDR kinases ATM and Chk2, multiple lysine/arginine acetylation and methylation events, phosphorylation by CK2 kinase on Ser392 in the CTD, and others^3^. PTMs control the recruitment of co-activators to p53 transactivation domains, trigger MDM2 disengagement to enhance protein stability and transactivation potential, and regulate DNA binding^3^.

*TP53* is the most frequently mutated gene in human and rodent cancers^12,13^. Interestingly, the overwhelming majority of these lesions are point mutations, some of which exert gain-of-function effects that, in addition to impairing tumor suppressor function, impart partial transforming activity to p53^14,15^. Recent studies have also shown a paradoxical pro-oncogenic role for wild-type p53 in certain contexts, such as in hepatocellular carcinoma^16^ and melanoma^17^. In HCC, *WT* p53 promotes a switch to cancer metabolism by suppressing oxidative phosphorylation and stimulating glycolysis. In melanoma, stress-induced Wnt5A signaling stabilizes p53 to elicit a slow cycling phenotype that facilitates resistance to therapy. Thus, p53 activity and function can be modulated in diverse ways to drive beneficial or adverse disease outcomes. These observations, as well as interest in pharmacological restoration of p53 tumor suppressor activity^18^, highlight the importance of gaining a full understanding of mechanisms that regulate p53 in tumor cells.

The pro-senescence activity of the transcription factor C/EBP*β* is inhibited in cancer cells by the 3’ untranslated region (3’UTR) of its mRNA^19,20^. To extend this novel mechanism to other proteins, we sought to determine if p53 is constrained in a similar manner. In this study we show that p53 is functionally suppressed in cancer cells by its 3’UTR. The 3’UTR effect involves subcellular partitioning of *TP53* transcripts away from a kinase-rich perinuclear cytoplasmic region, which prevents the encoded protein from accessing CK2, thus limiting phosphorylation on Ser392. In response to p53 activating signals such as oncogenic RAS in senescent cells and DNA damage in tumor cells, *TP53* mRNAs become perinuclear, coinciding with increased p53 phosphorylation and activation. Thus, *TP53* mRNA localization confers tight control over p53 tumor suppressor activity in proliferating and cancerous cells and is modulated by stress stimuli to facilitate p53 activation.

## Results

### The *TP53* 3’UTR decreases p53 cytostatic activity in human lung adenocarcinoma cells

To test whether the tumor suppressor activity of p53 is controlled by its 3’UTR, we generated Doxycycline (Doxy)-inducible lentiviral vectors containing human *TP53* cDNAs that either lack or include the 3’UTR (*TP53*^*CR*^ and *TP53*^*UTR*^, respectively). These constructs and the empty vector (Vec) were introduced into the p53-deficient human lung adenocarcinoma cell line, H1299, which carries an *NRAS*^*G12D*^ mutation. The cell lines were maintained as polyclonal pools. The two *TP53* constructs produced similar induced levels of p53 protein (Fig. 1a), indicating that the 3’UTR does not appreciably affect p53 expression, consistent with a recent report^21^. As expected, *TP53*^*CR*^ induction substantially reduced cell proliferation. *TP53*^*UTR*^ also suppressed proliferation but the effect was significantly less than for *TP53*^*CR*^ (Fig. 1a). Similar results were seen in colony formation assays (Fig. 1b). Expression of inducible murine *Trp53*^*CR*^ and *Trp53*^*UTR*^ constructs in NIH3T3 fibroblasts showed that the two proteins decreased proliferation of non-transformed cells equivalently (Supplementary Data Fig. 1a). By contrast, in HRAS^G12V^-tranformed cells *Trp53*^*CR*^ decreased proliferation while *Trp53*^*UTR*^ had little effect. Thus, the 3’UTR suppresses the RAS-dependent cytostatic activity of p53 without affecting protein levels.

**Fig. 1.**
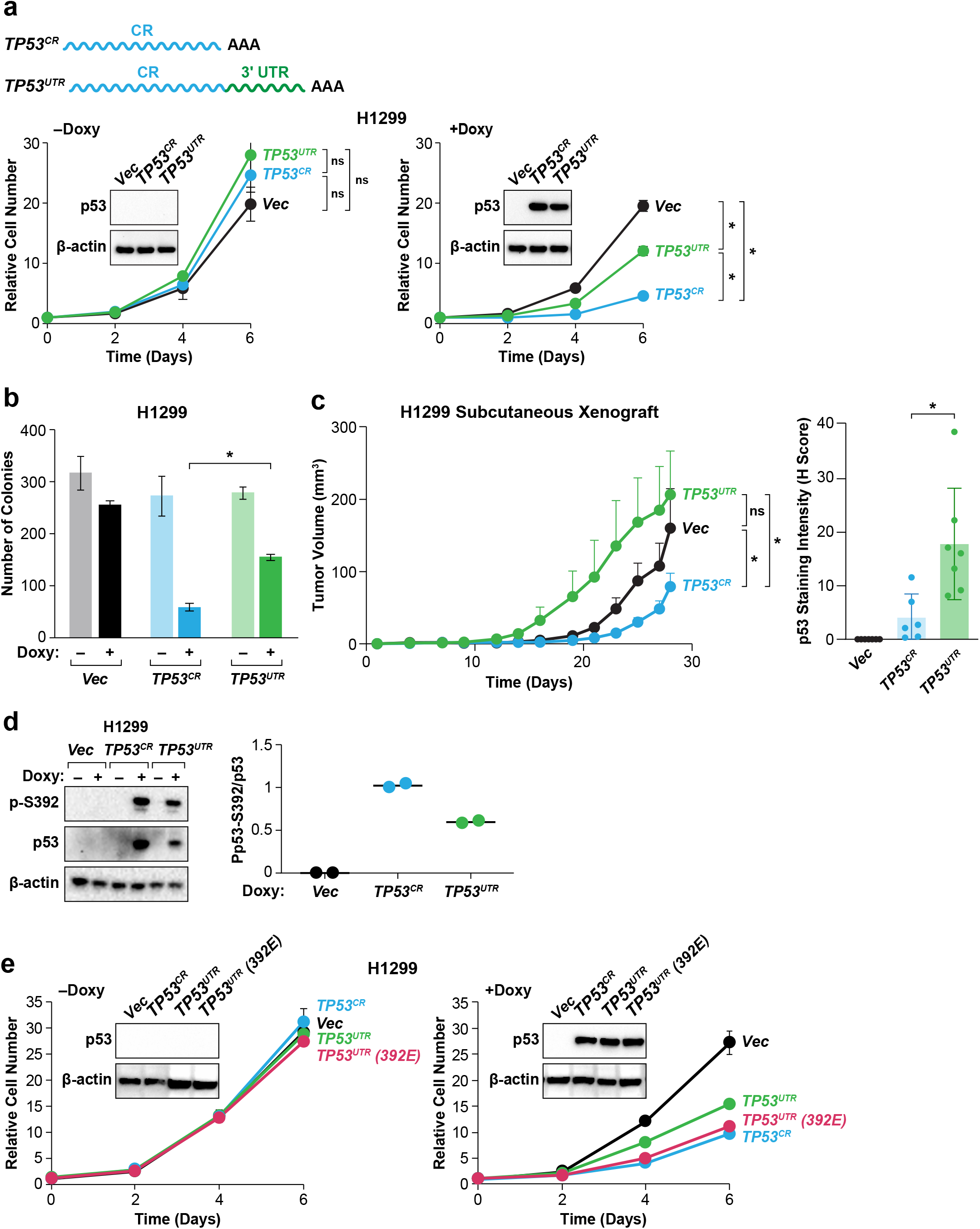
The *TP53* 3’UTR inhibits p53 tumor suppressor activity and phosphorylation on the Ser392 CK2 site in tumor cells. **a** Doxycycline-inducible constructs containing the human *TP53* coding region alone (*TP53*^*CR*^), coding region plus 3’UTR (*TP53*^*UTR*^), or empty vector (Vec) were introduced into *TP53* null human H1299 lung adenocarcinoma cells (*NRAS*^*Q61K*^). Cell proliferation was measured in the absence or presence of Doxycycline (Doxy). Data are the average of three independent cell populations for each construct. Immunoblots (insets) show equivalent p53^CR^ and p53^UTR^ protein expression. **b** Colony growth of p53-inducible H1299 cell lines ± Doxy. **c** H1299 cell lines were injected bilaterally into the flanks of athymic Nu/Nu mice and tumor growth was monitored under continuous feeding with Doxy diet. N=6 or 7 animals/group; data are the group averages ± SEM. p53 levels were determined by IHC staining of one tumor per mouse followed by quantitation of scanned images (H-scores) (right). **d** The 3’UTR suppresses p53 phosphorylation on a critical CK2 site, Ser392. Ser392 phosphorylation in p53^CR^ and p53^UTR^ proteins was analyzed by immunoblotting using an anti-p-Ser392 antibody. p-Ser392 levels in +Doxy samples, normalized to total p53, were quantified in two experiments (right). **e** A phosphomimetic S392E substitution overrides p53 inhibition by its 3’UTR. H1299 cells transduced with a Doxy-inducible *TP53*^*UTR*^ (S392E) construct were analyzed in cell proliferation assays. *TP53*^*CR*^ and *TP53*^*UTR*^ (± Doxy) were included for comparison.

To extend these findings to an *in vivo* setting, we transplanted the inducible *TP53*^*CR*^ and *TP53*^*UTR*^ H1299 cells subcutaneously into both flanks of athymic nude mice and monitored tumor growth. The experiment was performed under inducing conditions (Doxy diet) as we observed partial induction of the transgenes in mice on control diet. *TP53*^*CR*^ expression delayed tumor development relative to Vec control cells (Fig. 1c). However, tumors from *TP53*^*UTR*^-expressing cells grew equivalently or even faster than the controls, and significantly more than *TP53*^*CR*^ (P<0.05; two-way ANOVA). One tumor from each mouse was harvested at endpoint (26 days) and fixed sections were analyzed for p53 expression by IHC staining. Quantitation of scanned images (Fig. 1c, right panel) showed low but detectable levels of p53^CR^ but more than four-fold higher levels of p53^UTR^ (P<0.05; unpaired t-test with Welch’s correction). We infer that the much higher levels of p53^UTR^ are due to selection against p53^CR^-expressing cells during tumor growth. Overall, these data demonstrate that the *TP53* 3’UTR inhibits the tumor suppressor activity of p53 *in vitro* and *in vivo*.

p53 activity is stimulated by CK2-mediated phosphorylation on Ser392 (Ser389 in mouse), including causing increased DNA binding of recombinant p53^22,23^, and the phosphorylation-deficient S392A mutant displays impaired tumor suppressor activity^24^. CK2 is an oncogenic kinase^25^ that acts as a RAS pathway effector through its re-localization to the perinuclear cytoplasm in transformed cells^20^. We used a p53 anti-p-Ser392 antibody to assess whether this phospho-modification occurs in p53-expressing H1299 cells, where signaling is driven by the endogenous NRAS oncoprotein. As shown in Fig. 1d, p-Ser392 was evident in p53^CR^ expressed in H1299 cells and was reduced in p53^UTR^. This result was confirmed using mouse *Trp53* constructs transiently expressed in *TP53* null PC-3 cells (Supplementary Data Fig. 1b). These cells do not carry a RAS oncogene, which allowed us to evaluate the effect of co-expressed HRAS^G12V^ on p-Ser389 levels. Oncogenic HRAS increased p53^CR^ phosphorylation by 3.4-fold but only 1.9-fold for p53^UTR^, providing further evidence that the 3’UTR suppresses RAS-induced p53 phosphorylation by CK2.

To determine whether 3’UTR inhibition of p53 activity involves reduced Ser392/389 phosphorylation, we generated an inducible murine *Trp53*^*UTR*^ construct carrying a phospho-mimetic S389E substitution. This protein displayed cytostatic activity in H1299 cells that was indistinguishable from that of p53^CR^, overriding the effect of the 3’UTR (Fig. 1e). Thus, suppressed phosphorylation on Ser392/389 is a key mechanism by which the 3’UTR inhibits p53 tumor suppressor activity.

### Analysis of target genes preferentially activated by the p53^CR^-encoded protein

To identify target genes that may be differentially regulated by p53^CR^ and p53^UTR^, we performed RNA-seq on H1299 cells carrying the inducible constructs. RNA was prepared from cell replicates for each construct at 0 or 24 h post induction and subjected to total RNA sequencing. Statistical cutoffs of ≥1.5-fold change and FDR≤ 0.05 were used to define differentially-expressed genes (DEGs) that are up-or down-regulated by each p53 protein, subtracting any Doxy-induced genes in Vec cells. 760 genes showed altered expression (up or down) by p53^CR^, while 376 were controlled by p53^UTR^ (Fig. 2a). 273 genes were common to both lists, leaving 487 genes selectively regulated by p53^CR^ and 103 by p53^UTR^. We then sought to identify p53^CR^-upregulated DEGs that were preferentially induced by p53^CR^ vs. p53^UTR^ [i.e., genes with a CR/UTR fold-change ratio (FCR) ≥1.5], as these candidates may account for the elevated growth arrest and tumor suppression activity of p53^CR^. This analysis produced a set of 75 genes shown in the heat map of Fig. 2b, where genes are ranked by their FCR; note that these genes can be increased, decreased or unchanged by p53^UTR^. The majority of these are protein-coding genes.

**Fig. 2.**
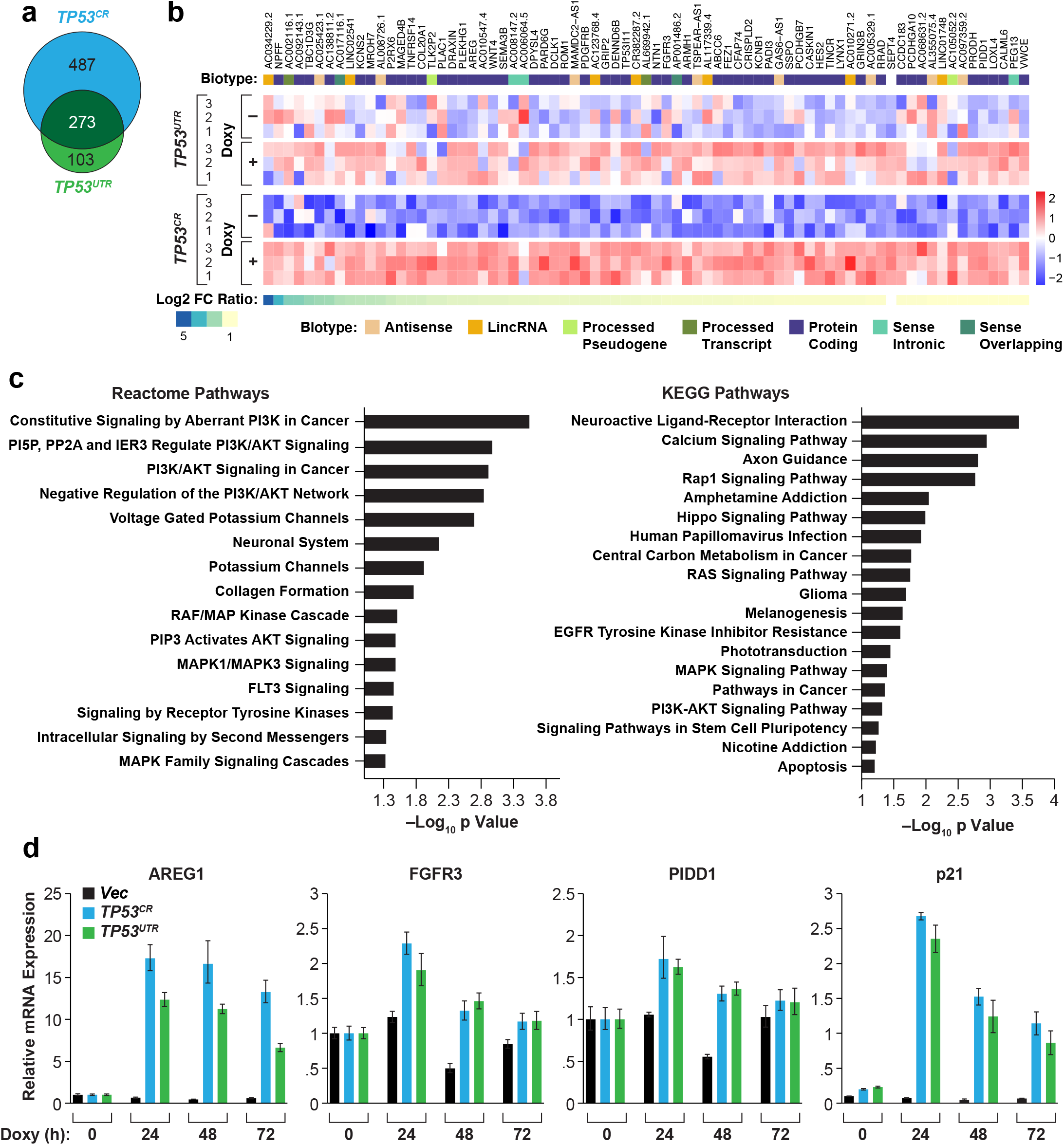
Effect of the 3’UTR on the p53-induced transcriptome. **a** Venn diagram of genes induced or down-regulated by *TP53*^*CR*^ or *TP53*^*UTR*^ in H1299 cells. The gene sets include transcripts that increase or decrease by ≥ 1.5-fold (FDR ≤ 0.05). **b** Heat map of the 75 *TP53*^*CR*^ up-regulated genes with a *TP53*^*CR*^:*TP53*^*UTR*^ fold-change ratio ≥ 1.5, sorted by log2(FC ratio). The heat map shows relative expression levels of each gene across the three *TP53*^*CR*^ and *TP53*^*UTR*^ replicates, ± Doxy. **c** Over-representation analysis of the differentially-regulated genes showing enrichment against KEGG and Reactome gene sets. **d** RT-qPCR analysis of pro-senescence/apoptotic candidate genes *AREG1, FCFR3* and *PIDD1* throughout an induction time course for H1299-*TP53*^*CR*^ and H1299-*TP53*^*UTR*^ cells. Transcript levels of the canonical p53 target, p21 (*CDKN1A*), are also shown. Data are averages from the three replicate cell lines.

The group of 75 genes was queried by over-representation analysis of KEGG and Reactome^26^ gene sets. The top scoring categories were skewed towards signaling pathways (Fig. 2c). Within these functional groups were potential p53 effectors associated with signaling such as *AREG* (senescence and apoptosis)^27,28^ and *FGFR3* (tumor suppression in epithelial cells)^29^, as well as genes involved in apoptosis (*PIDD1*^30^ and *SEPT-4*^31^). Each of these except *FGFR3* has been reported as a p53 target. RT-qPCR assays showed that *AREG, FGFR3* and *PIDD1* were at least modestly stimulated by p53^CR^ (Fig. 2d). However, preferential induction by p53^CR^ was seen only for *AREG*, which was also the most highly induced gene. The well-known p53 target and cell cycle arrest gene, *CDKN1A* (p21), was also induced equally by p53^CR^ and p53^UTR^ (Fig. 2d). Currently it is unclear whether a specific target gene is responsible for the selective cytostatic activity of p53 expressed without its 3’UTR or whether increased levels of many effectors cumulatively elicit this response.

### The 3’UTR excludes *TP53* transcripts from a perinuclear compartment containing CK2

The activity of the pro-senescence transcription factor C/EBP*β* is inhibited in tumor cells by 3’UTR-mediated exclusion of its mRNA from a perinuclear compartment enriched in the C/EBP*β* activating kinases, ERK1/2 and CK2^19,20^. To determine whether endogenous *TP53* transcripts show similar exclusion from the perinuclear cytoplasm, we performed combined RNA FISH and CK2*α* IF staining of RAS-transformed NIH3T3 cells and human tumor cell lines representing various cancer types that carry *WT* or mutant p53. In all cases, the perinuclear region containing CK2*α* was largely devoid of *TP53* mRNAs (Fig. 3a; Supplementary Fig. 2). Thus, mouse and human *TP53* transcripts partition away from the kinase-rich nuclear-proximal compartment in transformed cells.

**Fig. 3.**
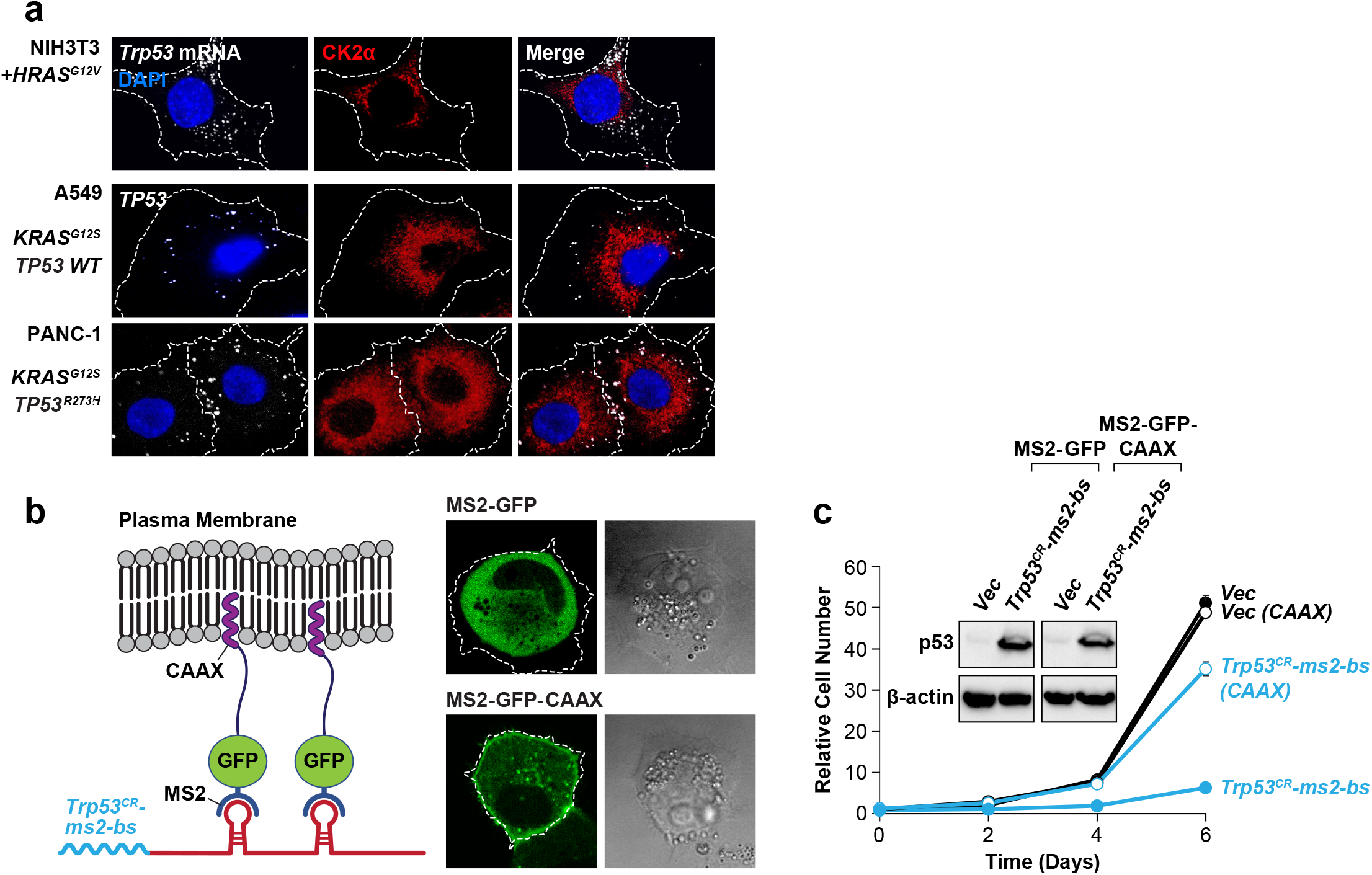
3’UTR-mediated exclusion of *TP53* transcripts from a perinuclear region enriched for CK2 suppresses p53 activity in tumor cells. **a** Localization of *TP53* mRNAs relative to CK2*α* in *RAS*-transformed tumor cells. Cells were analyzed by RNA FISH labeling of *TP53* transcripts together with immunostaining for CK2*α* and visualized by laser scanning confocal microscopy. Cell borders are indicated. **b** Strategy for mis-localizing *Trp53*^*CR*^ transcripts to the plasma membrane by MS2 tethering. MS2-GFP was fused to the *HRAS* CAAX membrane localization element and *Trp53*^*CR*^ was appended with multiple MS2 binding sites (*Trp53*^*CR*^-ms2-bs). Fluorescence images on the right show H1299 cells stably expressing MS2-GFP or MS2-GFP-CAAX. **c** Proliferation assays of MS2-GFP and MS2-GFP-CAAX cells infected with *Trp53*^*CR*^-ms2-bs or control lentiviral vectors. p53 levels (inset) are shown for the two transduced cell lines.

We next asked whether exclusion of *TP53* transcripts from the perinuclear CK2 domain is important for inhibition of p53 activity in transformed cells. To address this question, we applied a “ knock sideways” approach, using the MS2 RNA-binding protein to tether *Trp53*^*CR*^ mRNAs containing multiple MS2 binding sites (ms2-bs) to the plasma membrane (PM). The HRAS CAAX lipidation motif, which targets RAS to the PM, was appended to MS2-GFP^32^ (lacking the nls). The MS2-GFP-CAAX and MS2-GFP constructs were stably expressed in H1299 cells. Fluorescence imaging confirmed that MS2-GFP-CAAX was enriched at the PM, in contrast to more uniform localization of MS2-GFP (Fig. 3b). We then expressed *Trp53*^*CR*^-ms2-bs in MS2-GFP and MS2-GFP-CAAX cells and monitored cell proliferation. *Trp53*^*CR*^-ms2-bs suppressed proliferation of MS2-GFP cells as compared to the vector control (Fig. 3c). However, this cytostatic activity was largely absent in MS2-GFP-CAAX cells, despite equivalent p53 expression in the two cell lines. These results are consistent with the notion that *TP53* transcripts must partition to the perinuclear region for p53 to acquire tumor suppressor activity.

### Role of a U-rich element (URE) and the URE binding protein, HuR, in 3’UTR-dependent mRNA localization

The *TP53* 3’UTR contains a U-rich element (URE) that is a prominent feature of the mouse and human sequences^33^. The longer human 3’UTR also includes an ARE (A/U-rich element) located at its 3’ end. To assess whether the 3’UTR controls *Trp53* mRNA localization and the potential role of the URE in this process, we generated a mouse *Trp53*^*UTR*^ construct lacking the URE (*Δ*URE) and another in which the URE alone was fused to the coding region (*Trp53*^*URE*^) (Fig. 4a). *Trp53*^*CR*^, *Trp53*^*UTR*^ and the two mutant constructs were appended with ms2-bs repeats and expressed with the nuclear GFP-MS2-nls reporter^34^ in normal and RAS-transformed NIH3T3 cells. Fluorescence imaging showed that *Trp53*^*CR*^ mRNAs were distributed throughout the cytoplasm, while *Trp53*^*UTR*^ was more peripheral and was absent from the perinuclear space (Fig. 4b). *Trp53*^*Δ*^_*URE*_ transcripts were localized similarly to *Trp53*^*CR*^, while *Trp53*^*URE*^ mRNAs mirrored the distribution of *Trp53*^*UTR*^. These localization patterns were similar in transformed and non-transformed cells. The data suggest that the URE motif is necessary and sufficient to restrict *Trp53* transcripts away from the perinuclear region.

**Fig. 4.**
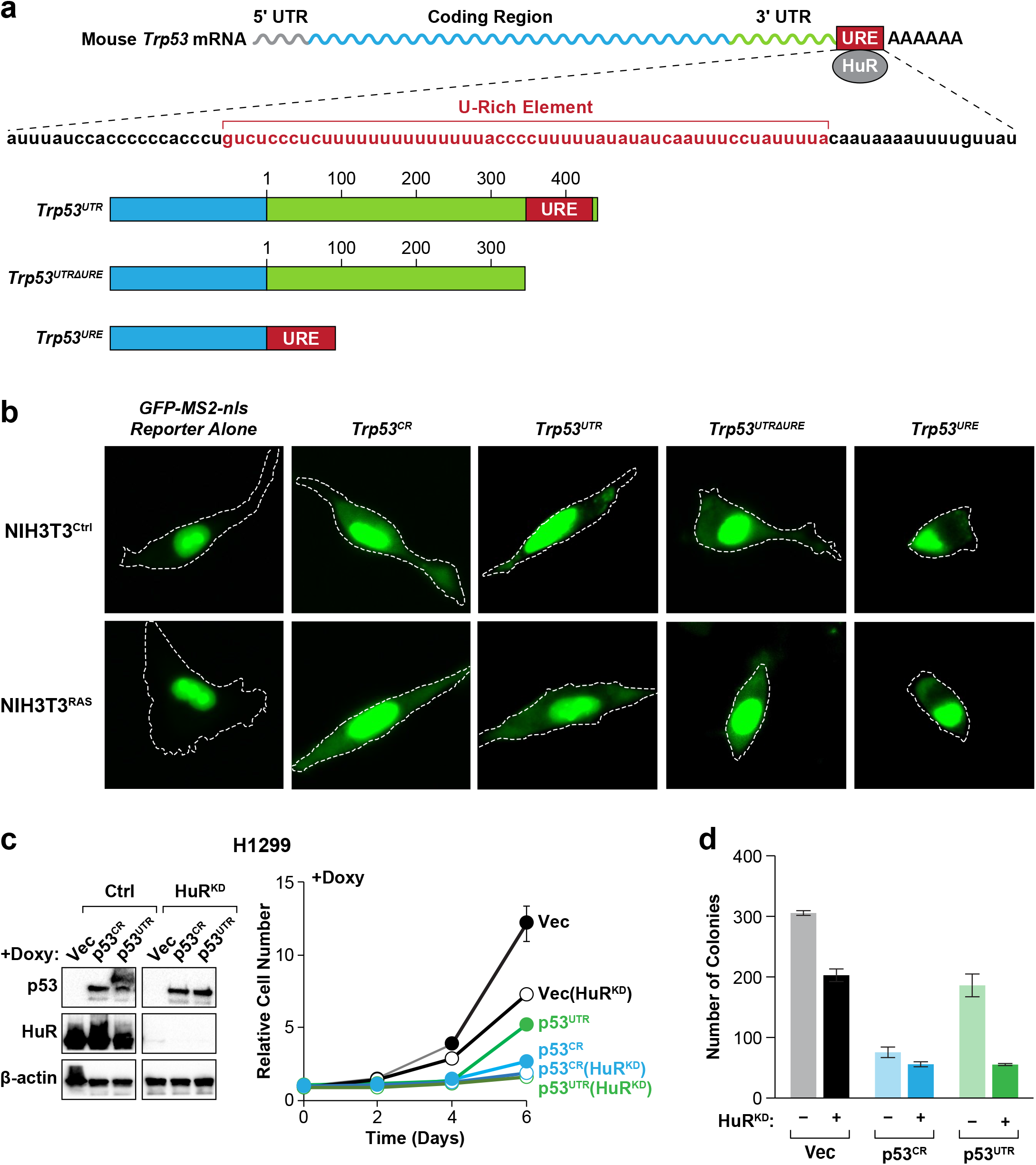
A U-rich element (URE) and its binding protein, HuR, are required for p53 3’UTR inhibition. **a** Location of a U-rich element (URE) in the 3’UTR of murine *Trp53*. Human *TP53* (not shown) contains a similar sequence in its 3’UTR^33^. Maps of the *Trp53* constructs used are shown (bottom). **b** MS2-GFP tagging of mouse *Trp53*^*CR*^, *Trp53*^*UTR*^, *Trp53*^*Δ*^_*URE*_ and *Trp53*^*URE*^ mRNAs reveals their cytoplasmic locations in control and *HRAS*-transformed NIH3T3 cells. Cells were transfected with the MS2-GFP-nls reporter alone or with the indicated constructs appended with MS2 binding sites. Representative live cell images are shown. Note the perinuclear exclusion of GFP signals for *Trp53*^*UTR*^ and *Trp53*^*URE*^. **c** Depletion of HuR abrogates the inhibitory effect of the *TP53* 3’UTR. *TP53*-inducible H1299 cell lines were infected with shHuR lentivirus or the empty vector and cell proliferation was analyzed in the presence of Doxy. p53 and HuR levels are shown on the left. **d** Colony formation assays using cells described in (**c)**.

The RNA-binding protein HuR/Elav1 was reported to bind the p53 URE motif^33^. HuR stabilizes many target mRNAs encoding proteins involved in cell proliferation and tumorigenesis, and high cytoplasmic HuR levels correlate with tumor malignancy^35,36^. We used native RNA immunoprecipitation (RIP) to confirm that endogenous *Trp53* transcripts associate with HuR in normal and RAS-transformed NIH3T3 cells (Supplementary Fig. 3a). We then asked whether HuR is required for the 3’UTR to inhibit p53 activity. HuR was depleted in the p53-inducible H1299 cell lines and proliferation was analyzed in the presence of Doxy (Fig. 4c). Loss of HuR did not affect p53 levels but abrogated the growth advantage of *TP53*^*UTR*^ cells compared to *TP53*^*CR*^. The proliferation rate of HuR-depleted *TP53*^*UTR*^ cells was similar to that of HuR-proficient *TP53*^*CR*^ cells, which grew poorly irrespective of HuR levels. Comparable effects were seen in clonogenic growth assays (Fig. 4d). Proliferation of Vec control cells also decreased modestly upon HuR depletion, as was expected considering the pro-oncogenic functions of HuR.

To evaluate the role of HuR in subcellular localization of endogenous *TP53* transcripts, we performed RNA FISH in transformed NIH3T3^RAS^ cells without and with HuR knockdown. HuR silencing caused marked perinuclear re-localization of *TP53* mRNA FISH signals, which were otherwise more peripherally distributed in cells expressing a control shRNA (Supplementary Fig. 3b). These and other data (see below) identify a critical role for HuR in controlling perinuclear exclusion of *TP53* transcripts, likely through binding to the URE motif.

### The p53 3’UTR inhibition mechanism is abrogated in senescent cells and requires AMPK*α*2

Since p53 is activated in primary cells undergoing OIS, we sought to determine if the 3’UTR inhibitory mechanism is nullified in these cells. We first confirmed that expression of oncogenic HRAS^G12V^ in MEFs increased p53 p-Ser389 levels compared to control cells, and this increase was blocked by the CK2 inhibitor CX-4945^37^ (Supplementary Fig. 4a). RAS-induced growth arrest was also completely abrogated by CX-4945, as the treated cells proliferated indistinguishably from MEFs without HRAS^G12V^ (Supplementary Fig. 4b). These results indicate that RAS-induced activation of p53 by CK2 plays a key role in OIS and suggest that p53 3’UTR inhibition is disabled in these cells. Accordingly, expression of either *Trp53*^*CR*^ or *Trp53*^*UTR*^ in HRAS^G12V^-transduced *p53*^*−/−*^ MEFs strongly inhibited colony formation (Fig. 5a).

**Fig. 5.**
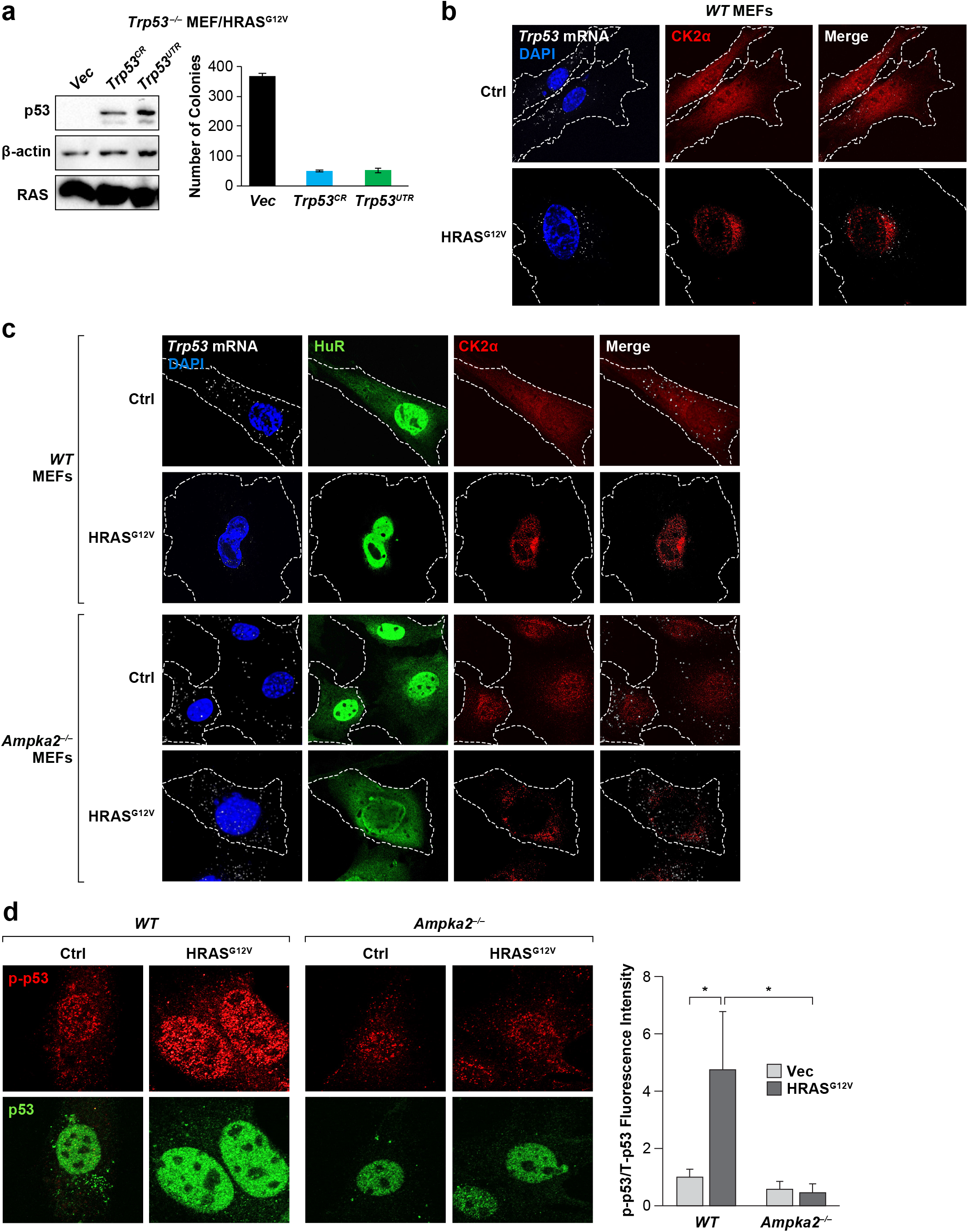
3’UTR-mediated p53 inhibition is abrogated in senescent primary MEFs undergoing *HRAS*^*G12V*^-induced senescence and requires AMPK*α*2. **a** *Trp53*^*CR*^ and *Trp53*^*UTR*^ suppress colony formation equally in *HRAS*^*G12V*^-transformed *Trp53*^*−/−*^ MEFs. Low passage *Trp53*^*−/−*^ MEFs expressing *HRAS*^*G12V*^ were infected with the two *Trp53* lentiviruses and colony growth was assayed. Equivalent p53 levels were expressed from each construct (left). **b** *HRAS*^*G12V*^ induces perinuclear clustering of *Trp53* transcripts and CK2*α* in MEFs. *WT* MEFs were infected with control and *HRAS*^*G12V*^ lentiviruses and analyzed for *Trp53* mRNA (RNA FISH) and CK2*α* (IF). **c** *HRAS*^*G12V*^-induced perinuclear re-localization of *Trp53* transcripts nuclear translocation of HuR in MEFs requires AMPK*α*2. *WT* and *Ampka2*^*−/−*^ MEFs were infected with control and *HRAS*^*G12V*^ lentiviruses and analyzed as described in (**b**), with the addition of HuR IF staining. **d** *WT* and *Ampka2*^*−/−*^ MEFs without or with *HRAS*^*G12V*^ were co-stained using antibodies for p-Ser389/392 and total p53. Nuclear signals were quantified using ImageJ and p-p53/total p53 ratios were calculated for each cell (right).

HuR undergoes nuclear translocation in senescent cells through an AMPK-dependent pathway, resulting in low cytoplasmic HuR^38,39,40^ which could inactivate the 3’UTR inhibitory mechanism. Therefore, we examined *Trp53* mRNA localization in *WT* MEFs before and after HRAS^G12V^ expression. RAS induced perinuclear clustering of *Trp53* transcripts which coincided with a more nuclear proximal distribution of CK2^20^ (Fig. 5b). We then asked whether this change in *TP53* mRNA localization requires AMPK activity. MEFs lacking the AMPK catalytic subunit, AMPK*α*2, but not AMPK*α*1, bypass OIS and continue to proliferate without increased p53 accumulation^41^. Therefore, we performed the same experiment in *Ampka2*^*−/−*^ MEFs. Cytoplasmic HuR levels remained elevated following HRAS^G12V^ expression in *Ampka2*^*−/−*^ cells and *Trp53* transcripts remained peripherally localized (Fig. 5c). Nevertheless, RAS-induced perinuclear translocation of CK2 still occurred, similar to *WT* cells. HRAS^G12V^ also increased p-Ser389 levels, normalized to total p53, in *WT* MEFs (Fig. 5d). However, p53 phosphorylation was abrogated in *Ampka2*^*−/−*^ MEFs, consistent with the more peripheral distribution of *Trp53* transcripts. These results indicate that cytoplasmic HuR is involved in shunting *Trp53* transcripts away from the perinuclear region. In OIS cells, AMPK*α*2 is required for nuclear transfer of HuR, allowing *Trp53* mRNAs to populate the perinuclear compartment which leads to p53 activation by kinases such as CK2.

### DNA damage induces perinuclear localization of *TP53* mRNA in tumor cells through AMPK*α*2 and ATM to promote p53 activation

p53 escapes activation by oncogenic signaling in tumor cells, although it remains responsive to AMPK agonists^42^. To determine whether the AMPK-p53 connection involves mRNA re-localization, we expressed a constitutively active form of AMPK^43^ in NIH3T3^RAS^ cells. CA-AMPK provoked distinct nuclear-proximal clustering of *Trp53* transcripts, coinciding with reduced cytoplasmic HuR levels (Supplementary Fig. 5a). Hence, AMPK signaling can induce *Trp53* mRNA translocation in transformed cells.

DNA damage stabilizes and activates p53 in tumor cells^44^ via the DDR and its effector kinases^45^. We investigated whether treatment of tumor cells with DNA damaging agents alters the peripheral localization of *TP53* transcripts. Human A549 lung adenocarcinoma cells (*WT TP53*; *KRAS*^*G12S*^) were exposed to 25 *μ*M cisplatin (CDDP), a DNA cross-linking agent, or vehicle for 16 h and cells were imaged for *TP53* mRNA and CK2*α* (Fig. 6a). *TP53* transcripts became more perinuclear following CDDP administration, and nuclear FISH signals were apparent in treated but not untreated cells, suggesting reduced nucleocytoplasmic mRNA export. CK2 was perinuclear in untreated cells and this was accentuated following DNA damage. HuR also displayed CDDP-induced nuclear translocation (Fig. 6b). As in senescent MEFs, altered *TP53* mRNA partitioning was associated with increased phosphorylation on Ser392 (p-p53/total p53; Fig. 6c). CCDP-treated cells also showed a large increase in p53 levels that can be attributed to its stabilization by DDR kinases, an effect that is less apparent in OIS MEFs (Fig. 5d).

**Fig. 6.**
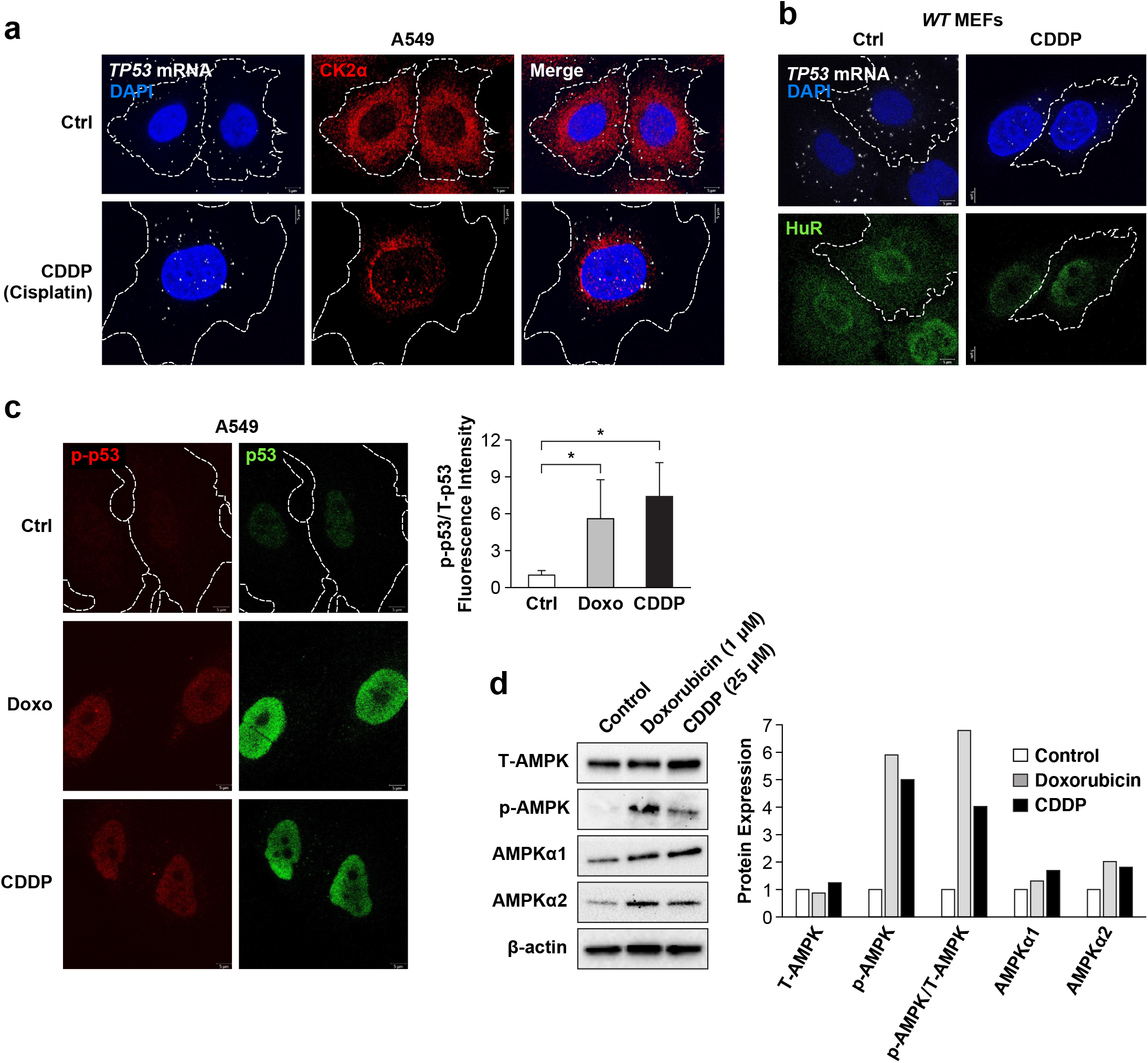
DNA damage induces perinuclear re-localization of *TP53* transcripts coinciding with HuR nuclear translocation, p53 phosphorylation and AMPK activation. **a** A549 cells were treated with 25 *μ*M cisplatin (CDDP) or vehicle for 16 h and analyzed for subcellular localization of *TP53* mRNA and CK2*α*. **b** A549 cells were treated as described in (**a**) and analyzed for subcellular localization of *TP53* mRNA and HuR. **c** DNA damage induces phosphorylation on p53 Ser392 in A549 cells. Cells were treated with 1 *μ*M doxorubicin or 25 *μ*M CDDP and co-immunostained for p-Ser392 and total p53. Normalized p-Ser392 levels (right) were quantitated from cell images. **d** DNA damage activates AMPK without significantly increasing AMPK*α*1/2 levels. A549 cells were treated with doxorubicin or CDDP for 16 h and lysates analyzed by immunoblotting for the indicated proteins. p-AMPK denotes p-Thr172 on AMPK*α*. Chemiluminescent signals were quantified and normalized to *β*-actin (right).

To address whether DNA damage-induced perinuclear localization of *TP53* transcripts is associated with AMPK activation, we treated A549 cells with the topoisomerase II inhibitor Doxorubicin (Doxo) or CCDP and assessed phosphorylation on AMPK*α* Thr172, the major phosphoacceptor for AMPK activation (Fig 6d). Both agents increased p-Thr172 by 5-to 6-fold but did not significantly enhance levels of AMPK*α*1, AMPK*α*2 or total AMPK*α*. Thus, DNA damage mainly augments AMPK activity rather than its levels.

We next used the AMPK inhibitor, Compound C, to confirm the functional role of AMPK activation in DNA damage-induced re-localization of *TP53* transcripts. A549 cells exposed to CDDP or Doxo displayed perinuclear *TP53* mRNA, concurrent with HuR nuclear uptake. Both of these responses were efficiently blocked by treating cells with 10 *μ*M Compound C. (Fig. 7a). DNA damage-induced phosphorylation on p53 Ser392 was also suppressed by the inhibitor (Fig. 7b). In addition, silencing of AMPK*α*2 prevented perinuclear segregation of *TP53* mRNAs and HuR translocation triggered by DNA damage (Fig. 7c), while AMPK*α*1 knockdown had no discernable effect (Supplementary Fig. 5b). Hence, genotoxic agents, acting specifically through AMPK*α*2, stimulate p53 activation in tumor cells through spatial reprogramming of *TP53* transcripts.

**Fig. 7.**
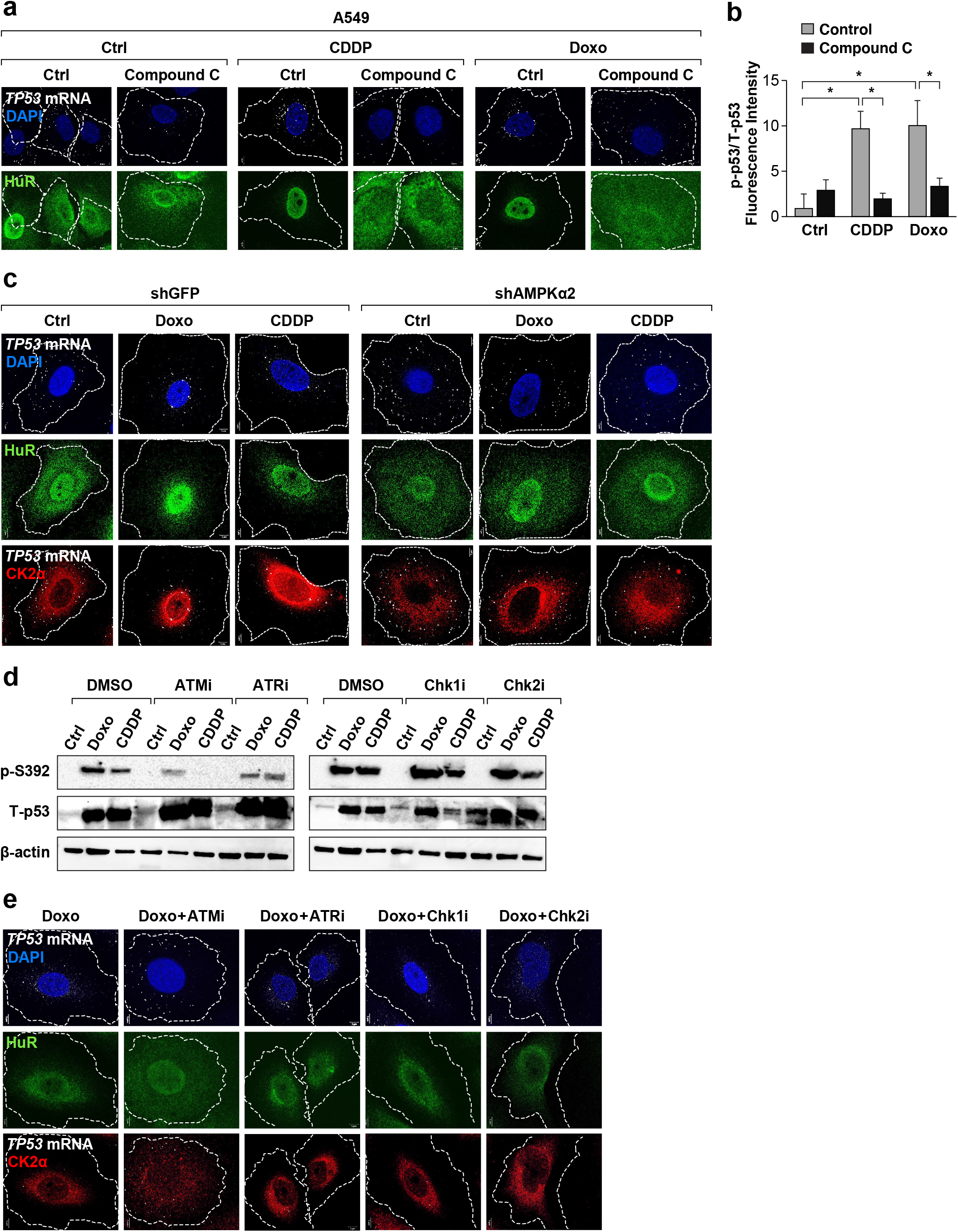
AMPK*α*2 and ATM mediate DNA damage-induced perinuclear translocation of *TP53* mRNA and p53 phosphorylation. **a** A549 cells were treated with CDDP or doxorubicin for 16 h in the absence or presence of the AMPK inhibitor, Compound C (10 *μ*M). Cells were analyzed for subcellular localization of *TP53* transcripts and HuR. **b** Normalized p-p53 (p-Ser392/total p53) levels in the cells described in (**a**), as determined by image analysis of co-stained cells. **c** Subcellular localization of *TP53* mRNA, HuR, and CK2*α* in A549 cells treated with cisplatin or doxorubicin for 16 h, ± AMPK*α*2 depletion. **d** DNA damage-induced phosphorylation on p53 Ser392 in the absence or presence of DDR kinase inhibitors. A549 cells were treated for 24 h with doxorubicin or CDDP, ± inhibitors for ATM (KU-60019), ATR (AZD6738), Chk1 (LY2603618) or Chk2 (BML277). Cell lysates were analyzed by immunoblotting for p-Ser392, total p53, and *β*-actin. **e** Doxorubicin-treated A549 cells were analyzed for subcellular localization of *TP53* mRNA, HuR, and CK2*α* in the absence or presence of DDR kinase inhibitors as described in (**d**).

DNA damage induces the ATM-Chk2 and ATR-Chk1 pathways through double-strand breaks or single-strand breaks/fork stalling, respectively, to activate p53 and other targets and elicit the appropriate cellular responses^46^. We used inhibitors of DDR kinases to investigate their potential roles in *TP53* mRNA re-localization following DNA damage, initially examining p53 Ser392 phosphorylation by immunoblotting (Fig. 7d). Only the ATM inhibitor (KU-60019) caused a significant decrease in p-Ser392 levels induced by either CDDP or Doxo, both of which can induce DSBs to activate ATM. These data suggest that DSB-ATM signaling may provoke perinuclear re-localization of *TP53* transcripts to facilitate p53 phosphorylation by CK2. Accordingly, ATMi alone was able to reverse the perinuclear partitioning of *TP53* transcripts in Doxo-treated A549 cells, re-establishing the peripheral, dispersed pattern seen in unstressed cells (Fig. 7e). Furthermore, the ATM inhibitor restored cytoplasmic HuR levels whereas the other inhibitors did not. ATMi also caused CK2 to become less perinuclear, although the significance of this response and whether it is an on-or off-target effect is presently unclear. These findings identify ATM as a key intermediate in the DDR signaling pathway that induces spatial reprogramming of *TP53* transcripts and phosphorylation/activation of p53.

### MDM2 is required for peripheral partitioning of *TP53* mRNAs in tumor cells

The MDM2 oncoprotein is a critical negative regulator of p53 in tumor cells. MDM2 is also a target of ATM, becoming phosphorylated on several sites in response to DNA damage that diminish MDM2 stability and oligomerization and decrease its activity toward p53^47,48,49,50^. In light of ATM’s role in redistribution of *TP53* mRNAs following DNA damage (Fig. 7d,e), we hypothesized that MDM2 may be involved in regulating peripheral localization of *TP53* transcripts in transformed/immortalized cells to suppress p53 activity. Supporting this idea, Mdm2 depletion in NIH3T3 cells triggered nuclear translocation of HuR and tight perinuclear clustering of *Trp53* mRNAs (Fig. 8a). Similar effects of Mdm2 silencing were observed in HRAS^G12V^-transformed *p19*^*Arf−/−*^ MEFs (Fig. 8b). In these cells, loss of Mdm2 also up-regulated p53 phosphorylation on Ser389 (Fig. 8c).

**Fig. 8.**
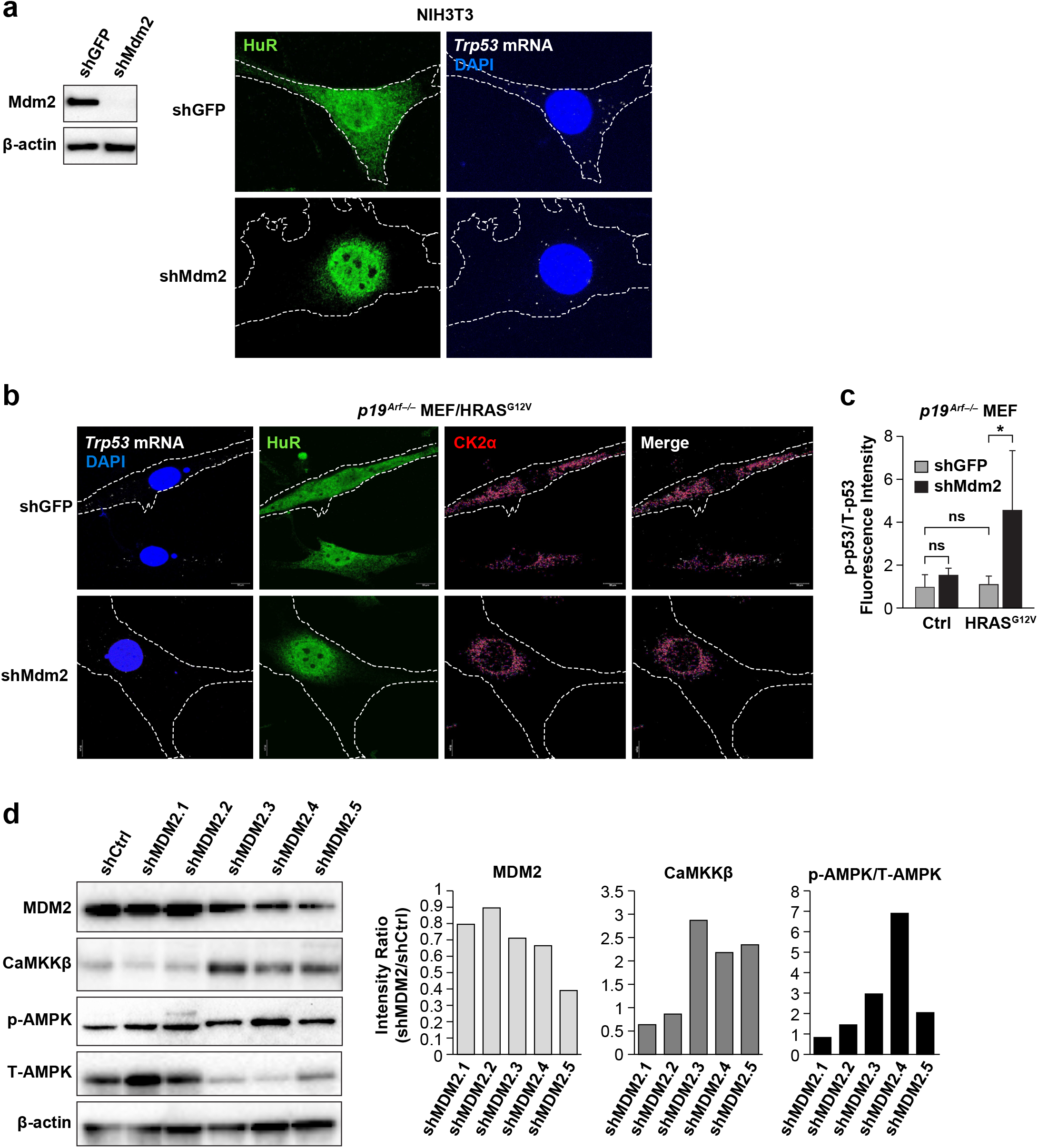
MDM2 enforces cytoplasmic HuR localization, peripheral compartmentalization of *TP53* transcripts and reduced p53 phosphorylation. **a** Mdm2 silencing in NIH3T3 cells induces nuclear translocation of HuR and increases nuclear proximity of *Trp53* transcripts. Mdm2 knockdown was confirmed by immunoblotting (left). **b** Mdm2 depletion in *HRAS*^*G12V*^-transformed *p19*^*Arf−/−*^ MEFs increases perinuclear clustering of *Trp53* mRNAs and overlap with CK2*α*, along with HuR nuclear translocation. **c** Mdm2 silencing in *HRAS*^*G12V*^-transformed *p19*^*Arf−/−*^ MEFs stimulates p53 phosphorylation on Ser389. Normalized p53 phosphorylation was determined by co-staining cells with antibodies for p-Ser389 and total p53 and measuring fluorescence intensity ratios. **d** MDM2 knockdown in A549 cells increases levels of the AMPK kinase, CaMKK*β*, and increased phosphorylation on AMPK*α* Thr172. Five different MDM2 shRNA constructs were tested; efficient MDM2 silencing correlates with increased CaMKK*β* expression. Band intensities were quantitated from gel scans (right).

To extend these observations, we silenced MDM2 in A549 tumor cells using five independent shRNAs and examined AMPK signaling. For the three shRNAs that produced the most efficient MDM2 knockdown, the ratio of p-AMPK to total AMPK was increased (Fig. 8d). This result suggests that MDM2 may be involved in suppressing AMPK signaling in tumor cells, consistent with the peripheral localization of *TP53* transcripts. AMPK activation through AMPK*α* Thr172 phosphorylation is catalyzed mainly by the upstream kinases LKB1 (STK11) or CaMKK*β*^51^. Since A549 cells carry a mutation in the *STK11* tumor suppressor gene and thus lack LKB1^52^, another kinase must be responsible for activating AMPK. Therefore, we analyzed CaMKK*β* levels in the knockdown cells, also since this kinase was implicated in AMPK*α*2 activation in cells undergoing RAS-induced senescence^53^. CaMKK*β* expression increased substantially in cells with efficient MDM2 knockdown (Fig. 8d). These results suggest that MDM2 is involved, directly or indirectly, in maintaining low CaMKK*β* levels in transformed cells to inhibit the AMPK-p53 axis by ensuring peripheral localization of *TP53* mRNAs.

## Discussion

We show here that the *TP53* 3’UTR restrains post-translational activation of p53 in tumor cells. Our findings provide an important addition to the repertoire of mechanisms that negatively regulate p53 activity in tumors and unstressed normal cells. This additional layer of control provides further protection from inappropriate activation of p53, which can be toxic at the cellular and organismal levels^7^. Our previous studies showed that the pro-senescence functions of C/EBP*β* are also suppressed by its 3’UTR, a process termed 3’UTR regulation of protein activity (UPA)^19,53^. p53 UPA shares many of the same properties, particularly the key role of mRNA localization and the requirement for HuR, as well as reversal of UPA by AMPK signaling. It is anticipated that other proteins with tumor suppressor functions will be modulated by similar or related 3’UTR-dependent mechanisms.

AMPK*α*2 is a key regulator for perinuclear re-localization of *TP53* transcripts and the associated activation of p53 induced by genotoxic or oncogenic stresses. AMPK*α*2 is likewise required for C/EBP*β* activation during OIS, also by abrogating the 3’UTR inhibitory mechanism^53^. These and other findings^41^ implicate AMPK*α*2 as part of a critical upstream pathway involving CaMKK*β* that unleashes the anti-oncogenic activities of p53 and C/EBP*β* and therefore must be disabled in cancer cells. Studies are currently underway in our laboratory to investigate the putative tumor suppressor function of AMPK*α*2 using mouse models. Importantly, our work also identifies MDM2 as a negative regulator of the CaMKK*β*-AMPK*α*2-p53 axis. The fact that MDM2 enforces peripheral localization of *TP53* mRNAs could account for much of its p53-restraining activity in tumor cells.

The effect of AMPK signaling on p53 activation is indirect, as it promotes p53 phosphorylation by CK2 (Figs. 5d and 7b,d). AMPK was reported to phosphorylate p53 on Ser15 in response to glucose deprivation^43^. However, the Ser15 motif does not resemble a classical AMPK phosphoacceptor site and is thought to be a target of DDR kinases, DNA-PK, and/or p38^3^. Whether Ser15 phosphorylation and other p53 modifications are subject to 3’UTR regulation requires further investigation. At present it is unknown how many p53 kinases besides CK2 and ERK1/2^20^ are perinuclear in transformed and senescent cells. Such information could provide clues to the extent of p53 regulation by its 3’UTR, as spatial partitioning between kinases and mRNAs is critical to the UPA mechanism.

H1299 cells expressing *TP53*^*CR*^ displayed reduced proliferation and an enlarged cell morphology characteristic of senescence, but were not positive for the canonical senescence marker, SA-*β*Gal. This can be explained by the fact that C/EBP*β* is also inhibited by UPA in tumor cells, and C/EBP*β* is required for SA-*β*Gal positivity in senescing cells^53^. p53 and C/EBP*β* appear to be in parallel UPA pathways, such that expression of the active p53^CR^ protein does not override C/EBP*β* 3’UTR inhibition. Thus, C/EBP*β* remains functionally repressed in tumor cells forced to express active p53, whereas upstream stimuli such as metabolic stress or AMPK agonists (e.g., metformin), or oncogenic RAS in primary cells^41,53,54^, can activate both proteins to elicit a comprehensive senescent response.

Suppression of p53 Ser392/389 phosphorylation by CK2 appears to be an important target of 3’UTR repression, since mutating this position to a phosphomimetic residue overcame 3’UTR-mediated suppression of p53 cytostatic activity. Ser392 phosphorylation is induced by DNA damage^22^ and regulates both *WT* and oncogenic forms of p53. An alanine substitution at Ser392 increased the transforming activity of p53 mutant R175H in combination with oncogenic RAS; conversely, cells expressing p53 R175H combined with phosphomimetic S392E were more sensitive to the cytotoxic effects of genotoxic agents^24^. The same study found low levels of Ser392 phosphorylation in human breast tumors expressing mutant p53, which is consistent with our finding that both *WT* and mutant *TP53* transcripts segregate away from the perinuclear CK2 region in tumor cells (Fig. 3a and Supplementary Fig. 2). The reversal of p53 3’UTR inhibition by DNA damaging agents may explain why tumor cells with mutant p53 retain some sensitivity to chemotherapy. Further characterization of the p53 UPA machinery may uncover novel targets for adjuvant drugs that potentiate tumor cell responses to chemotherapeutic agents.

## Acknowledgements

We thank Donna Butcher (MHL, Laboratory Animal Sciences Program, Leidos Biomedical Research, Inc., Frederick National Laboratory for Cancer Research) for performing IHC staining of tumor samples and Allen Kane (Scientific Publications, Graphics & Media, Leidos Biomedical Research, Inc., Frederick National Laboratory for Cancer Research) for preparation of figures. This research was supported in part by the Intramural Research Program of the NIH, National Cancer Institute, Center for Cancer Research. This work utilized the computational resources of the NIH HPC Biowulf cluster (http://hpc.nih.gov). The content of this publication does not necessarily reflect the views or policies of the Department of Health and Human Services, nor does mention of trade names, commercial products, or organizations imply endorsement by the U.S. Government.

## Methods

### Mice and preparation of MEFs

Mice were maintained in accordance with National Institutes of Health animal guidelines following protocols approved by the NCI-Frederick Animal Care and Use Committee. *p19*^*Arf−/−*^ (*CdKn2a-Arf*) mice were obtained from the NCI-Frederick repository on a mixed background (B6.129), backcrossed to C57Bl/6Ncr and maintained as a homozygous stock. *Ampka2*^*−/−*^ (C57Bl/6Ncr) mice have been described^53,55^. *TP53*^*−/−*^ mice (kindly provided by Dr. Glenn Merlino) were maintained as heterozygotes backcrossed to C57Bl/6Ncr, and matings between heterozygous carriers were performed to generate mouse embryonic fibroblasts (MEFs). MEFs were isolated from *WT, p19*^*Arf−/−*^, *Ampka2*^*−/−*^ and *Trp53*^*−/−*^ embryos at E13.5. The cells were maintained at low passage.

### Cells and cell culture

H1299, A375, RKO, PANC-1, HepG2, HEK-293T and GP2-293 cells were cultured in DMEM medium (Gibco) supplemented with 10% FBS (Gibco) and 100 μg/ml normocin (InvivoGen, ant-nr-1). A549 cells were cultured in F-12K Medium (Gibco) with 10% FBS and 100 μg/ml normocin. MEFs were cultured in DMEM with 10% FBS and 100 μg/ml primocin (InvivoGen, ant-pm-05), IMR90 cells were purchased from ATCC and cultured in EMEM (Gibco) with 10% FBS and 100 μg/ml primocin. NIH3T3 cells were cultured in DMEM with 10% calf serum (Colorado Serum Company) and 100 μg/ml normocin. MIA PaCa-2 cells were purchased from ATCC and cultured in DMEM with 10% FBS, 2.5% horse serum (Gibco) and 100 μg/ml normocin. Cell stocks were generally passaged fewer than 5 times, and freshly thawed cells were maintained in culture for no more than 2 weeks before conducting experiments.

### Antibodies and reagents

p53 (#2524) and phospho-p53 (p-Ser392; #9281) antibodies were from Cell Signaling Technology. Antibodies for *β*-actin (C11), CK2*α* (E-7), HuR (3A2), CaMKKβ (H-95), AMPKα1 (H-4; #SC-398861) and AMPKα2 (A-20 #SC-19129) were from Santa Cruz Biotechnology. p-AMPKα (#2531) and total-AMPKα (D5A2) antibodies were from Cell Signaling Technology. panRAS antibody (#610002) was from BD Biosciences. Antibodies for mouse Mdm2 (ab10344) and human MDM2 (ab38618) were from Abcam. Anti-mouse (W4028) and anti-rabbit (W4018) HPR-conjugated secondary antibodies were from Promega. Anti-Mouse Alexa Fluor 488 (A32790), anti-rabbit Alexa Fluor 594 (A21207) and Anti-Mouse Alexa Fluor 647 (A21236) secondary antibodies were from ThermoFisher Scientific. Doxorubicin and CDDP (Sigma), Doxycycline (Sigma-Aldrich #D9891), CX-4945 (Activate Scientific), and inhibitors from APExBIO for ATM (KU-60019; #A8336), ATR (AZD6738; #B6007), Chk1 (LY2603618; #A8638) and Chk2 (BML277; #B1236) were reconstituted according to the manufacturers’ recommendations and used at the indicated doses.

### Plasmids and lentiviral vectors

Murine and human *TP53* coding region (*TP53*^*CR*^) and coding region plus 3’UTR (*TP53*^*UTR*^) fragments were PCR amplified and inserted into pcDNA3.1. The fragments were then transferred to pBabe-puro and the Doxycycline-inducible lentiviral vector pINDUCER10b^56^. The GFP-MS2-nls reporter system has been described^34^. Multiple MS2 binding sites were inserted into mouse *Trp53* constructs using the Gateway multisite cloning system^57,58^ as follows: Four segments were used to insert a CMV51p promoter (attL4/L5), mouse *Trp53*^*CR*^ (attR5/R1), either MS2b or a stuffer sequence (attL1/L2), and finally mouse *Trp53 3’UTR, ΔURE, URE* or stuffer fragments (attR2/R3) to make sequence-verified entry clones that were incorporated into the attR4/R3 destination vector pDest667 (lentiviral vector with puro selection). Some of the resulting expression cassettes were moved to pBabe puro or pcDNA 3.1 using introduced flanking restriction sites and traditional cloning methods, as needed. Human HRAS^G12V^ expression vectors and lentiviral shRNA vectors targeting mouse and human AMPKα1, AMPKα2 and HuR have been described^53^. shRNA knockdown vectors for mouse Mdm2 (TRNC0000302276) or human MDM2 (TRCN0000355726, TRCN0000355727, TRCN0000355728, TRCN0000003378 and TRCN0000003380) were from Sigma Aldrich. The pBabe-puro expression vector for CA-AMPK has been described^53^.

### Retrovirus and lentivirus packaging

The retroviral packaging plasmid VSVG was co-transfected with retroviral plasmids into GP2-293 cells. A third-generation lentivirus packaging system consisting of 6 μg pMD2.G, 10 μg pMDLg/pRRE and 5 μg pRSV/Rev, together with 20 μg lentiviral plasmid, was used to transfect 293T cells. Transfection was performed by the CaPO_4_ precipitation method. After 48-72 h, viral supernatants were collected and passed through 0.45 *μ*m filters. Target cells were infected with viral supernatants in the presence of 8 *μ*g/mL Polybrene. After 48 h, cells were selected with the appropriate antibiotics.

### Xenograft experiments

Six-to eight-week-old Balb/c athymic nude mice were randomly grouped and injected subcutaneously (both flanks) with 1 × 10^6^ H1299 *TP53*-inducible cells suspended in 100 µl of phosphate buffer saline (PBS). At two days post injection, the mice were placed on 200 mg/kg Doxycycline diet (Bio-Serv, #S3888) daily. Tumor volume was monitored three times weekly by caliper measurements and was calculated as π/6 × length × width × height. Mice were euthanized at 26 days post-injection (in one case a tumor reached 2 cm prior to the scheduled endpoint and the animal was euthanized). Tumors were collected, fixed in 10% NBF for five days and then placed into 70% ethanol. Tissues were embedded and sectioned for subsequent analysis.

### IHC staining of tumor samples

Slides were generated from paraffin blocks using 5 *μ*M sections and stained with H&E following standard protocols. H&E staining was performed using the Sakura® Tissue-Tek® Prisma™ automated stainer. The slides were dewaxed using Xylene and then hydrated using a series of graded ethyl alcohols. Commercial Hematoxylin, Clarifier, Bluing Reagent and Eosin-Y were used to stain the slides. A regressive staining method was used, which intentionally overstains tissues and then uses a differentiation step (Clarifier/Bluing reagents) to remove excess stain. After staining was completed, the slides were cover-slipped using the Sakura® Tissue-Tek™Glass® automatic cover-slipper and dried. IHC staining was performed using LeicaBiosystems BondRX autostainer with the following conditions: Epitope Retrieval 1 (Citrate), 20 min; p53 Ab (Agilent #M7001, 1:25 dilution), 30 min; M.O.M.® (Mouse on Mouse) ImmPRESS® HRP Polymer Kit (Vector Labs #MP-2400); Bond Polymer Refine Detection Kit (LeicaBiosystems #DS9800) with omission of the PostPrimary Reagent and Polymer reagents. Mouse IgG2b Isotype Control (BD Biosciences) was used in place of p53 for the negative control. Slides were removed from the Bond autostainer, dehydrated through ethanols, cleared with xylenes, and cover-slipped. IHC slides were scanned at 20X using an Aperio AT2 scanner (Leica Biosystems, Buffalo Grove, IL) into whole slide digital images. IHC images were analyzed using cytonuclear algorithm in HALO imaging analysis software (v3.3.2541.300; Indica Labs, Corrales, NM) to determine percent positive cells and H score.

### Immunoblots

Cells were lysed in RIPA buffer and protein concentrations were determined using Bradford reagent (Bio-Rad Laboratories). Equal amounts of protein were applied to SDS-PAGE gels which were run at 150 V initially and 200 V thereafter. Proteins were then transferred to nitrocellulose membranes. Membranes were blocked with 4% milk in TBST for 1h at room temperature and incubated with primary antibodies at 4°C overnight. Membranes were washed with TBST three times, incubated with HRP-conjugated secondary antibodies at room temperature for 1 h and visualized using enhanced chemiluminescence substrate (SuperSignal™ West Dura Extended Duration Substrate #34076, Thermo Scientific).

### RNA sequencing and data analysis

Three biological replicates of stably transduced H1299-Vec, H1299-*TP53*^*CR*^, or H1299-*TP53*^*UTR*^ cells (1.5 × 10^6^) were seeded in 100 mm dishes for next day treatment with 0.1 *μ*g/ml Doxycycline or control (-Doxy). Cells were harvested 24 h later and RNA was isolated using the Gene Jet RNA Purification kit (Thermo Fisher). RNA was quantified and 10 *μ*g of each sample was treated with 2 units of DNase plus RNase inhibitor (Invitrogen) in 1X DNase buffer and nuclease free water. The reaction was incubated for 30 min at 37^°^C followed by mixing with two volumes of chloroform and incubating on ice for 5 minutes, centrifuged for 10 minutes at 12000 rpm (4^°^C). The upper layer was transferred and precipitated by adding two volumes of 100% ethanol and 1/10 volume of 3M sodium acetate at -20^°^C for 2 h. The mixture was centrifuged for 20 min at 14000 rpm (4^°^C) and RNA pellets were washed with 70% ethanol, resuspended in RNase free water, and quantified on nanodrop. The purified RNA samples were used for RNA sequencing.

The quality of the isolated RNA was validated prior to sequencing using an Ailgent Bioanalyzer. All samples had an RNA integrity number (RIN) greater than 9. Libraries were constructed for each p53 cell line using Illumina’s TruSeq mRNA Prep Kit (#RS-122–2101) following manufacturer’s instructions. Samples were multiplexed and sequenced on a NovaSeq 6000. The demultiplexed samples yielded between 45-61 million paired-end reads.

The quality of each sequenced sample was assessed using FastQC (version 0.11.5) [1], Preseq (version 2.0.3)^59^, Picard tools (version 1.119) (https://broadinstitute.github.io/picard/), FastQ Screen (0.9.3)^60^, Kraken (1.1)^61^, QualiMap (2.2.1)^62^, and RSeQC (version 2.6.4)^63^. Reads were trimmed using Cutadapt (version 1.18)^64^ to remove synthetic adapters sequences and low-quality bases. The trimmed reads were aligned against the human reference genome, hg38, using STAR (version 2.7.0f)^65^ in two-pass mode. The percentage of uniquely mapped reads was greater than 80.4% in all samples. Expression levels were quantified using RSEM (version 1.3.0)^66^ with GENCODE annotation version 30^67^. Low count genes were removed prior to differential expression analysis. Genes with a counts per million (CPM) greater than 0.5 in at least two samples were considered for downstream analysis.

The expected counts from RSEM were then normalized using the voom algorithm^62^ from the Limma R package (version 3.40.6)^68,69^. These normalized counts were used for clustering and data visualization. Limma was used to test for differential gene expression between the induced and control conditions: “ TP53^CR^ – Ctrl_TP53^CR^”, “ TP53^UTR^ – Ctrl_ TP53^UTR^” and “ Vec – Ctrl_Vec.” Significant differentially expressed genes (DEGs) were identified with an absolute fold-change ≥ 1.5 and a false-discovery rate ≤ 0.05. Any significant differentially-expressed genes found in the empty vector “ Vec – Ctrl_Vec” comparison were removed from the “ TP53^CR^ – Ctrl_TP53^CR^” and “ TP53^UTR^ – Ctrl_ TP53^UTR^” DEG lists. The resulting DEG lists were further categorized based on the direction of the fold-change to distinguish between up- and down-regulated genes to create the following groups: TP53^CR^_up, TP53^CR^_down, TP53^UTR^_up, and TP53^UTR^_down. To compare the results of the TP53^CR^ and TP53^UTR^ experimental conditions, a fold-change ratio was calculated for each gene in TP53^CR^_up, TP53^CR^_down, TP53^UTR^_up, and TP53^UTR^_down. Genes with an absolute fold-change ratio ≥ 1.5 were identified as differentially regulated between contrasts. TP53^CR^_up genes with fold-change ratios (FCRs) vs. TP53^UTR^ ≥ 1.5 were taken as TP53^CR^-specific genes and were analyzed further. Enriched pathways were identified by over-representation analysis of KEGG and Reactome^26^ gene sets from the Molecular Signatures Database^70^.

### RT-qPCR analysis of differentially expressed genes

H1299-Vec, H1299-*TP53*^*CR*^, and H1299-*TP53*^*UTR*^ cells were seeded (3 × 10^5^ cells/60 mm dish) and the next day induced with 0.1 *μ*g/ml Doxy and harvested at 0, 24, 48, and 72 h. RNA was isolated using the GeneJet RNA Purification kit (ThermoFisher Scientific) and cDNA was synthesized using Reverse Transcription Kit (Qiagen). Quantitative real time PCR was performed using SYBR Green with QuantiTect Primers (Qiagen) and using ppia gene as a reference gene. The primers were: Hs_AREG (Qiagen; #QT00030772); Hs_PIDD1 (Qiagen; #QT02406691); Hs_FGFR3 (Qiagen; #QT01000685); p21 (Bio-Rad; #qHsaCID0014498); Hs_PPIA (Qiagen; #QT01866137).

### Cell proliferation assays

2 × 10^4^ cells per well were seeded into 6-well plates. At each time point, cells were fixed with 10% formaldehyde in PBS for 15 min and stained with 0.1% crystal violet for 30 minutes. The dye was extracted using 10% acetic acid and absorbance was measured at 590 nm. All values were normalized to day 0 (24 h after plating).

### Colony formation assays

2 ×10^3^ H1299 cells were seeded into 10 cm plates and media was replaced every three days. After 10 days in culture, colonies were fixed with 10% formaldehyde for 15 minutes, stained with 0.1% crystal violet for 30 minutes and counted.

### GFP-MS2-nls RNA imaging

H1299, NIH3T3 and MEF cells were transfected using FuGENE6 (Qiagen), Polyfect (Qiagen), or X-treme GENE HP (Roche) reagents according to the manufacturer’s protocol. 5 × 10^4^ cells were seeded into glass bottom 35-mm dishes. After 16 h, cells were transfected with 0.2 *μ*g GFP-MS2-nls reporter and 2.3 *μ*g pcDNA-*Trp53*^*CR*^-MS2b or pcDNA-*Trp53*^*UTR*^-MS2b. 16 h after transfection, GFP positive cells were observed, and fluorescence images were acquired using a Zeiss LSM-780 confocal microscope.

### Immunofluorescence imaging

Cells were seeded on μ-Slides VI0.4 (Ibidi, 80606) in 100 *μ*l medium and fixed with 4% paraformaldehyde for 15 min and incubated with p53 antibody and p-p53 Ser389 (Ser392) antibody overnight at 4°C. Fluorescence images were acquired using a Zeiss LSM-780 confocal microscope.

### RNA fluorescence in situ hybridization (FISH)

Cells grown on μ-Slides VI0.4 (Ibidi 80606) were washed in freshly made CB buffer (10 mM MES pH 6.1, 150 mM NaCl, 5 mM MgCl_2_, 5 mM EGTA, 5 mM glucose). Cells were permeabilized and incubated in Pre-Fixative mix (2% paraformaldehyde, 0.01% glutaraldehyde, 0.05% saponin, in CB buffer) for 15 min at 4°C, then fixed with ice-cold Fixative mix (2% paraformaldehyde, 0.01% glutaraldehyde, in CB) for 15 min, rinsed in CB buffer twice at 4°C and quenched in 50 mM NH_4_Cl for 5 min at room temperature. RNA FISH procedure was carried out using probe hybridization, signal amplification and post-hybridization washes (QuantiGene ViewRNA ISH Cell Assay; Affymetrix, ThermoFisher) following the manufacturer’s instructions. The FISH probes were from ThermoFisher: mouse *Trp53* Type 4 488 probe (VA4-3082421-VC) ; human *TP53* Type 6 647 (VA6-13337-VC). For subsequent IF staining, cells were then rinsed with PBS-S (0.05% Saponin, in PBS) and blocked with 5% BSA in PBS-S for 1 h at room temperature and incubated with primary antibodies: CK2α (E-7), HuR (3A2), or CaMKKβ (H-95), overnight at 4°C. Cells were the rinsed three times for 5 min at room temperature with PBS-S and incubated with secondary antibodies for 1 h. DAPI was added at 0.1 mg/mL for 1 minute at room temperature. Stained cells were imaged using a Zeiss LSM-780 confocal microscope.

### RNA immunoprecipitation

NIH3T3 and NIH3T3^HRAS-G12V^ cells were washed twice with ice-cold PBS and lysed with LB buffer (137 mM NaCl; 20 mM Tris pH 8.0; 10% glycerol; 1% NP-40; 1 mM DTT; 1X protease inhibitor and RNase inhibitor) for 10 min on ice. The protein levels were determined using Bradford reagent (Bio-Rad Laboratories) and equalized. 10% of lysate was used for input after protein equalization. 1:50 HuR antibody was added and incubated for 3 h at 4°C. Protein G Sepharose 4 Fast Flow Beads (Millipore Sigma, GE17-0618-01) were added and incubated for 1 h at 4°C with rotation. RNA was prepared using the GeneJet RNA Purification kit (Thermo Fisher). First strand cDNA was synthesized using Reverse Transcription Kit (Qiagen). Quantitative real time PCR was performed using SYBR Green and the following primers from Qiagen: Ppia (QT00247709), Trp53(QT00101906). *Ppia* was used as a reference gene.

### Statistical analysis

Quantitative data are presented as means ± SD or SEM. Statistical analysis was performed in GraphPad Prism using Student’s t test or 2-way ANOVA functions. P-values less than 0.05 were considered significant and are indicated by an asterisk in the figures.

## Notes

### Competing Interest Statement

The authors have declared no competing interest.

## References

1. He, S. & Sharpless, N. E. Senescence in Health and Disease. Cell 169, 1000–1011 (2017).

2. Weber, J. D., Taylor, L. J., Roussel, M. F., Sherr, C. J. & Bar-Sagi, D. Nucleolar Arf sequesters Mdm2 and activates p53. Nat Cell Biol 1, 20–26 (1999).

3. Kruse, J. P. & Gu, W. Modes of p53 regulation. Cell 137, 609–622 (2009).

4. Vousden, K. H. & Prives, C. Blinded by the Light: The Growing Complexity of p53. Cell 137, 413–431 (2009).

5. Kubbutat, M. H., Jones, S. N. & Vousden, K. H. Regulation of p53 stability by Mdm2. Nature 387, 299–303 (1997).

6. Tollini, L. A., Jin, A., Park, J. & Zhang, Y. Regulation of p53 by Mdm2 E3 ligase function is dispensable in embryogenesis and development, but essential in response to DNA damage. Cancer Cell 26, 235–247 (2014).

7. Jones, S. N., Roe, A. E., Donehower, L. A. & Bradley, A. Rescue of embryonic lethality in Mdm2-deficient mice by absence of p53. Nature 378, 206–208 (1995).

8. Montes de Oca Luna, R., Wagner, D. S. & Lozano, G. Rescue of early embryonic lethality in mdm2-deficient mice by deletion of p53. Nature 378, 203–206 (1995).

9. Laptenko, O., et al. The p53 C terminus controls site-specific DNA binding and promotes structural changes within the central DNA binding domain. Mol Cell 57, 1034–1046 (2015).

10. Kim, E., Rohaly, G., Heinrichs, S., Gimnopoulos, D., Meissner, H. & Deppert, W. Influence of promoter DNA topology on sequence-specific DNA binding and transactivation by tumor suppressor p53. Oncogene 18, 7310–7318 (1999).

11. Cain, C., Miller, S., Ahn, J. & Prives, C. The N terminus of p53 regulates its dissociation from DNA. J Biol Chem 275, 39944–39953 (2000).

12. Kaiser, A. M. & Attardi, L. D. Deconstructing networks of p53-mediated tumor suppression in vivo. Cell Death Differ 25, 93–103 (2018).

13. Kastenhuber, E. R. & Lowe, S. W. Putting p53 in Context. Cell 170, 1062–1078 (2017).

14. Oren, M. & Rotter, V. Mutant p53 gain-of-function in cancer. Cold Spring Harb Perspect Biol 2, a001107 (2010).

15. Brosh, R. & Rotter, V. When mutants gain new powers: news from the mutant p53 field. Nat Rev Cancer 9, 701–713 (2009).

16. Kim, J., et al. Wild-Type p53 Promotes Cancer Metabolic Switch by Inducing PUMA-Dependent Suppression of Oxidative Phosphorylation. Cancer Cell 35, 191–203 e198 (2019).

17. Webster, M. R., et al. Paradoxical Role for Wild-Type p53 in Driving Therapy Resistance in Melanoma. Mol Cell 77, 681 (2020).

18. Khoo, K. H., Verma, C. S. & Lane, D. P. Drugging the p53 pathway: understanding the route to clinical efficacy. Nat Rev Drug Discov 13, 217–236 (2014).

19. Basu, S. K., et al. 3’UTR elements inhibit Ras-induced C/EBPbeta post-translational activation and senescence in tumour cells. EMBO J 30, 3714–3728 (2011).

20. Basu, S. K., et al. Oncogenic RAS-Induced Perinuclear Signaling Complexes Requiring KSR1 Regulate Signal Transmission to Downstream Targets. Cancer Research 78, 891–908 (2018).

21. Mitschka, S. & Mayr, C. Endogenous p53 expression in human and mouse is not regulated by its 3’UTR. eLife 10, (2021).

22. Keller, D. M., et al. A DNA damage-induced p53 serine 392 kinase complex contains CK2, hSpt16, and SSRP1. Mol Cell 7, 283–292 (2001).

23. Cox, M. L. & Meek, D. W. Phosphorylation of serine 392 in p53 is a common and integral event during p53 induction by diverse stimuli. Cell Signal 22, 564–571 (2010).

24. Yap, D. B., et al. Ser392 phosphorylation regulates the oncogenic function of mutant p53. Cancer Research 64, 4749–4754 (2004).

25. Roffey, S. E. & Litchfield, D. W. CK2 Regulation: Perspectives in 2021. Biomedicines 9, (2021).

26. Fabregat, A., et al. The Reactome Pathway Knowledgebase. Nucleic Acids Res 46, D649–D655 (2018).

27. Pommer, M., Kuphal, S. & Bosserhoff, A. K. Amphiregulin Regulates Melanocytic Senescence. Cells 10, (2021).

28. Taira, N., et al. Induction of amphiregulin by p53 promotes apoptosis via control of microRNA biogenesis in response to DNA damage. Proc Natl Acad Sci USA 111, 717–722 (2014).

29. Lafitte, M., et al. FGFR3 has tumor suppressor properties in cells with epithelial phenotype. Molecular Cancer 12, 83 (2013).

30. Lin, Y., Ma, W. & Benchimol, S. Pidd, a new death-domain-containing protein, is induced by p53 and promotes apoptosis. Nat Genet 26, 122–127 (2000).

31. Garcia-Fernandez, M., et al. Sept4/ARTS is required for stem cell apoptosis and tumor suppression. Genes Dev 24, 2282–2293 (2010).

32. Yan, X., Hoek, T. A., Vale, R. D. & Tanenbaum, M. E. Dynamics of Translation of Single mRNA Molecules In Vivo. Cell 165, 976–989 (2016).

33. Mazan-Mamczarz, K., et al. RNA-binding protein HuR enhances p53 translation in response to ultraviolet light irradiation. Proc Natl Acad Sci USA 100, 8354–8359 (2003).

34. Rook, M. S., Lu, M. & Kosik, K. S. CaMKIIalpha 3’ untranslated region-directed mRNA translocation in living neurons: visualization by GFP linkage. J Neurosci 20, 6385–6393 (2000).

35. Lopez de Silanes, I., Lal, A. & Gorospe, M. HuR: post-transcriptional paths to malignancy. RNA Biol 2, 11–13 (2005).

36. Abdelmohsen, K. & Gorospe, M. Posttranscriptional regulation of cancer traits by HuR. Wires Rna 1, 214–229 (2010).

37. Siddiqui-Jain, A., et al. CX-4945, an orally bioavailable selective inhibitor of protein kinase CK2, inhibits prosurvival and angiogenic signaling and exhibits antitumor efficacy. Cancer Research 70, 10288–10298 (2010).

38. Wang, W., et al. AMP-activated kinase regulates cytoplasmic HuR. Mol Cell Biol 22, 3425–3436 (2002).

39. Wang, W., et al. AMP-activated protein kinase-regulated phosphorylation and acetylation of importin alpha1: involvement in the nuclear import of RNA-binding protein HuR. J Biol Chem 279, 48376–48388 (2004).

40. Wang, W., Yang, X., Lopez de Silanes, I., Carling, D. & Gorospe, M. Increased AMP:ATP ratio and AMP-activated protein kinase activity during cellular senescence linked to reduced HuR function. J Biol Chem 278, 27016–27023 (2003).

41. Phoenix, K. N., Devarakonda, C. V., Fox, M. M., Stevens, L. E. & Claffey, K. P. AMPKalpha2 Suppresses Murine Embryonic Fibroblast Transformation and Tumorigenesis. Genes & cancer 3, 51–62 (2012).

42. Motoshima, H., Goldstein, B. J., Igata, M. & Araki, E. AMPK and cell proliferation--AMPK as a therapeutic target for atherosclerosis and cancer. J Physiol 574, 63–71 (2006).

43. Jones, R. G., et al. AMP-activated protein kinase induces a p53-dependent metabolic checkpoint. Mol Cell 18, 283–293 (2005).

44. Chen, X., Ko, L. J., Jayaraman, L. & Prives, C. p53 levels, functional domains, and DNA damage determine the extent of the apoptotic response of tumor cells. Genes Dev 10, 2438–2451 (1996).

45. Reinhardt, H. C. & Schumacher, B. The p53 network: cellular and systemic DNA damage responses in aging and cancer. Trends Genet 28, 128–136 (2012).

46. Awasthi, P., Foiani, M. & Kumar, A. ATM and ATR signaling at a glance. J Cell Sci 128, 4255–4262 (2015).

47. Cheng, Q., Chen, L., Li, Z., Lane, W. S. & Chen, J. ATM activates p53 by regulating MDM2 oligomerization and E3 processivity. EMBO J 28, 3857–3867 (2009).

48. Carr, M. I., Roderick, J. E., Gannon, H. S., Kelliher, M. A. & Jones, S. N. Mdm2 Phosphorylation Regulates Its Stability and Has Contrasting Effects on Oncogene and Radiation-Induced Tumorigenesis. Cell Rep 16, 2618–2629 (2016).

49. Gannon, H. S., Woda, B. A. & Jones, S. N. ATM phosphorylation of Mdm2 Ser394 regulates the amplitude and duration of the DNA damage response in mice. Cancer Cell 21, 668–679 (2012).

50. Cheng, Q. & Chen, J. Mechanism of p53 stabilization by ATM after DNA damage. Cell Cycle 9, 472–478 (2010).

51. Hardie, D. G., Schaffer, B. E. & Brunet, A. AMPK: An Energy-Sensing Pathway with Multiple Inputs and Outputs. Trends Cell Biol 26, 190–201 (2016).

52. Ji, H., et al. LKB1 modulates lung cancer differentiation and metastasis. Nature 448, 807–810 (2007).

53. Basu, S. K., et al. A RAS-CaMKKbeta-AMPKalpha2 pathway promotes senescence by licensing post-translational activation of C/EBPbeta through a novel 3’UTR mechanism. Oncogene 37, 3528–3548 (2018).

54. Serrano, M., Lin, A. W., McCurrach, M. E., Beach, D. & Lowe, S. W. Oncogenic ras provokes premature cell senescence associated with accumulation of p53 and p16INK4a. Cell 88, 593–602 (1997).

55. Laderoute, K. R., et al. 5’-AMP-activated protein kinase (AMPK) is induced by low-oxygen and glucose deprivation conditions found in solid-tumor microenvironments. Mol Cell Biol 26, 5336–5347 (2006).

56. Meerbrey, K. L., et al. The pINDUCER lentiviral toolkit for inducible RNA interference in vitro and in vivo. Proc Natl Acad Sci USA 108, 3665–3670 (2011).

57. Hartley, J. L., Temple, G. F. & Brasch, M. A. DNA cloning using in vitro site-specific recombination. Genome Res 10, 1788–1795 (2000).

58. Cheo, D. L., Titus, S. A., Byrd, D. R., Hartley, J. L., Temple, G. F. & Brasch, M. A. Concerted assembly and cloning of multiple DNA segments using in vitro site-specific recombination: functional analysis of multi-segment expression clones. Genome Res 14, 2111–2120 (2004).

59. Daley, T. & Smith, A. D. Predicting the molecular complexity of sequencing libraries. Nat Methods 10, 325–327 (2013).

60. Wingett, S. W. & Andrews, S. FastQ Screen: A tool for multi-genome mapping and quality control. F1000Res 7, 1338 (2018).

61. Wood, D. E. & Salzberg, S. L. Kraken: ultrafast metagenomic sequence classification using exact alignments. Genome Biol 15, R46 (2014).

62. Okonechnikov, K., Conesa, A. & Garcia-Alcalde, F. Qualimap 2: advanced multi-sample quality control for high-throughput sequencing data. Bioinformatics 32, 292–294 (2016).

63. Wang, L., Wang, S. & Li, W. RSeQC: quality control of RNA-seq experiments. Bioinformatics 28, 2184–2185 (2012).

64. Martin, M. Cutadapt removes adapter sequences from high-throughput sequencing reads. In: EMBnet (ed^(eds) (2011).

65. Dobin, A., et al. STAR: ultrafast universal RNA-seq aligner. Bioinformatics 29, 15–21 (2013).

66. Li, B. & Dewey, C. N. RSEM: accurate transcript quantification from RNA-Seq data with or without a reference genome. BMC Bioinformatics 12, 323 (2011).

67. Harrow, J., et al. GENCODE: the reference human genome annotation for The ENCODE Project. Genome Res 22, 1760–1774 (2012).

68. Law, C. W., Chen, Y., Shi, W. & Smyth, G. K. voom: Precision weights unlock linear model analysis tools for RNA-seq read counts. Genome Biol 15, R29 (2014).

69. Smyth, G. K. Linear models and empirical bayes methods for assessing differential expression in microarray experiments. Stat Appl Genet Mol Biol 3, Article3 (2004).

70. Liberzon, A., Subramanian, A., Pinchback, R., Thorvaldsdottir, H., Tamayo, P. & Mesirov, J. P. Molecular signatures database (MSigDB) 3.0. Bioinformatics 27, 1739–1740 (2011).

