## Supplementary Data for "3’UTR-dependent dynamic changes in *TP53* mRNA localization regulate p53 tumor suppressor activity"

### Supplementary Figure 1

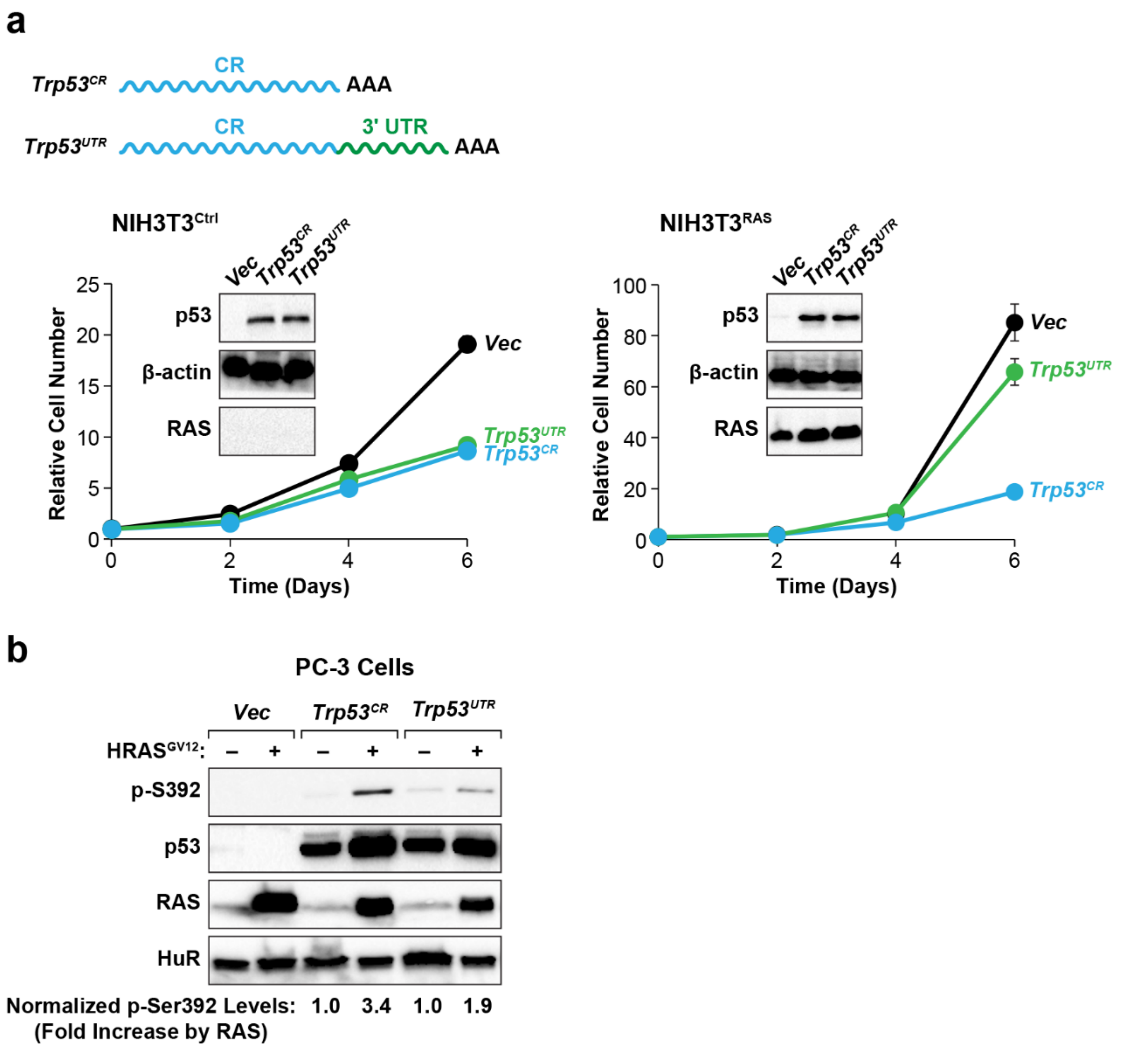


**Supplementary Fig. 1 The cytostatic activity of murine p53 is inhibited by its 3'UTR in *HRAS$^{G12V}$*-transformed NIH3T3 cells. a** *Trp53$^{CR}$* and *Trp53$^{UTR}$* lentiviruses and the empty vector were used to infect NIH3T3 control and HRAS$^{G12V}$-expressing cells (NIH3T3$^{RAS}$). Cell proliferation was analyzed over a six-day time course. Equivalent p53 expression is shown by immunoblots (insets). **b** HRAS$^{G12V}$-induced phosphorylation on p53 Ser392 is inhibited by the 3'UTR. Mouse *Trp53* constructs were transfected into PC-3 cells (*TP53* null, *PTEN* null human prostate cancer) without and with expression of HRAS$^{G12V}$. Cell lysates were analyzed by immunoblotting for p-Ser392, total p53 and RAS. HuR was used as a loading control. Normalized p-S392 levels are the ratio of p-S392 to total p53 band intensities and are expressed as fold increase by HRAS$^{G12V}$.

### Supplementary Figure 2

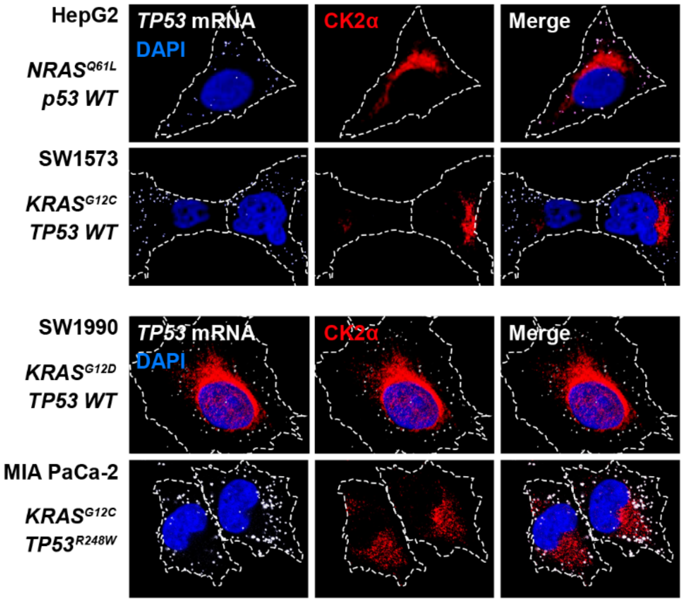


**Supplementary Fig. 2** ***WT*** **and mutant** ***TP53*** **transcripts are excluded from a perinuclear CK2 compartment in human tumor cell lines. a** HepG2 (hepatocarcinoma), SW1573 (lung adenocarcinoma), SW1990 (metastatic pancreatic adenocarcinoma), and MIA PaCa-2 (pancreatic ductal adenocarcinoma) cell lines were analyzed by RNA FISH for *TP53* mRNA and IF staining for CK2α. The cells were stained with DAPI to visualize nuclei. *RAS* and *TP53* mutant status for each cell line is indicated.

### Supplementary Figure 3

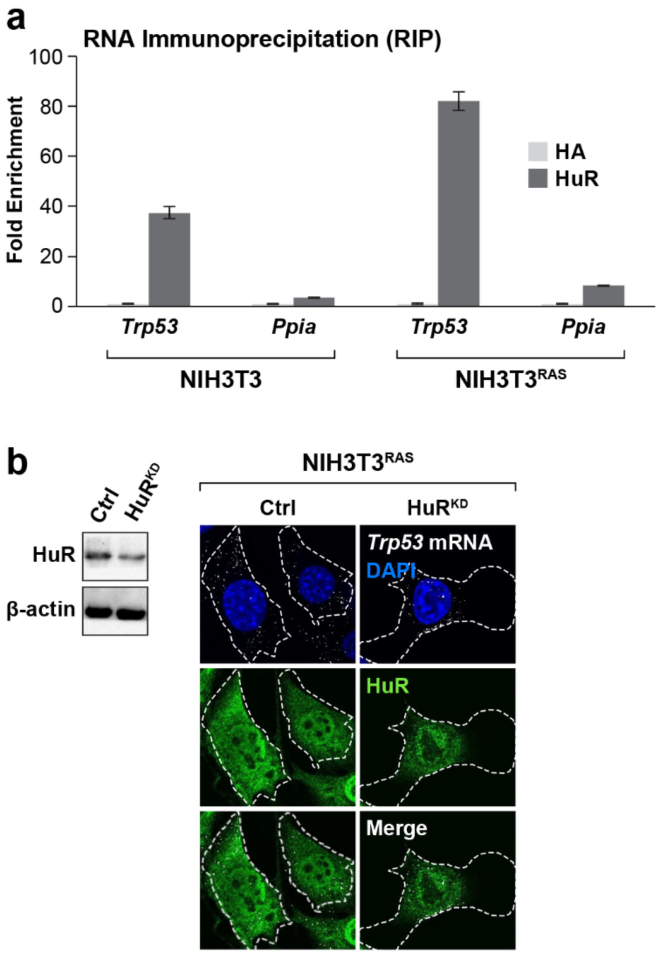


**Supplementary Fig. 3 HuR binds to *Trp53* transcripts and regulates mRNA localization. a** Native RNA immunoprecipitation (RIP) was performed on NIH3T3 and NIH3T3$^{RAS}$ cell lysates using antibodies for HA (negative control) and HuR. Input and RIP RNA samples were analyzed by RT-qPCR using primers for *Trp53* and *Ppia* (non-specific reference mRNA). RIP:input mRNA ratios were calculated to determine fold enrichment. **b** HuR depletion causes perinuclear translocation of *Trp53* transcripts. NIH3T3$^{RAS}$ cells were infected with shHuR and control shRNA lentiviruses and analyzed by RNA FISH to detect *Trp53* transcripts and immunostained for HuR. HuR knockdown was also confirmed by immunoblotting (left).

**Supplementary Figure 4**

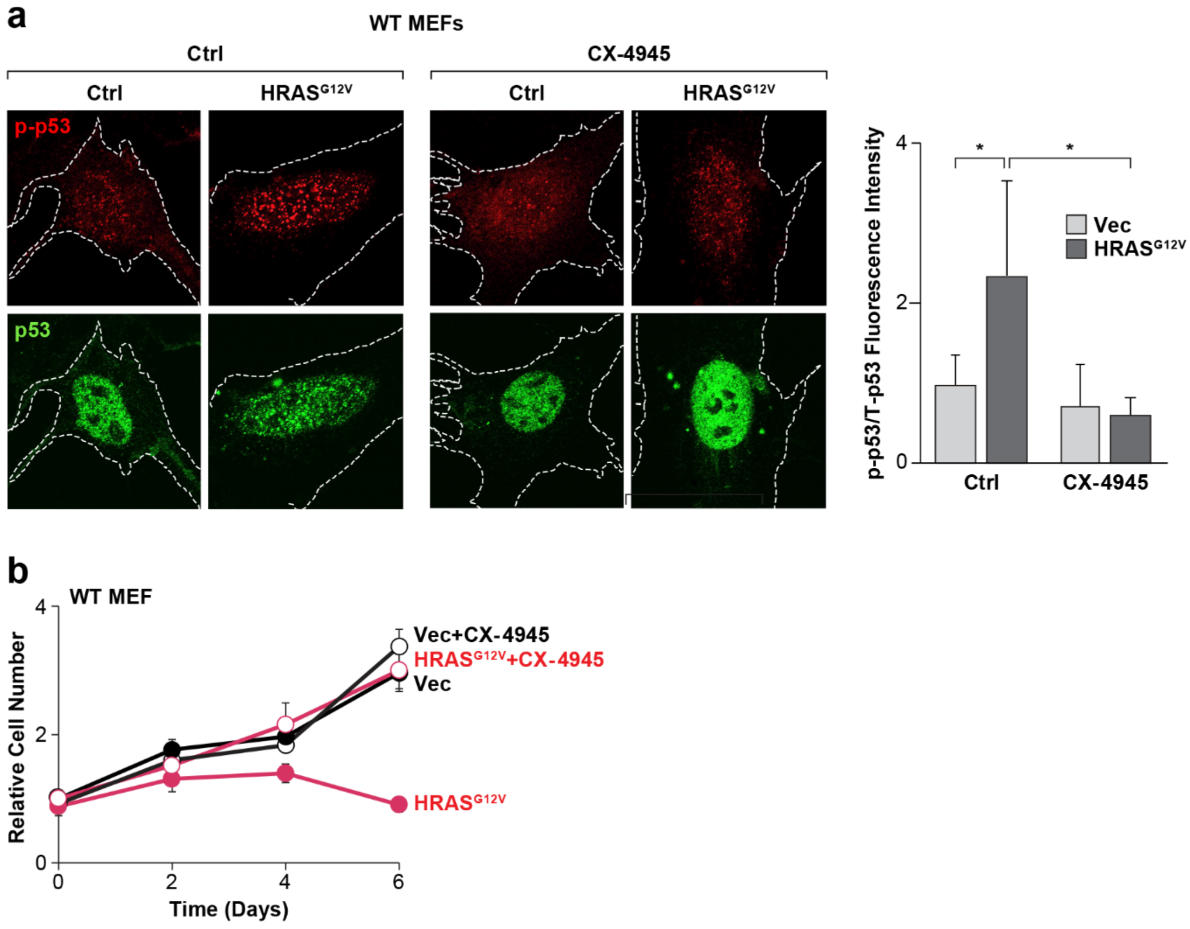


**Supplementary Fig. 4 CK2 inhibition blocks *HRAS$^{G12V}$*-induced phosphorylation on Ser389 and senescence-associated growth arrest in MEFs. a** MEFs were infected with control and *HRAS$^{G12V}$* vectors, treated without or with 5 μM CX-4945 (CK2 inhibitor) for 16 h and co-immunostained for p-Ser389 and total p53. Fluorescence intensity ratios were determined per cell (right). **b** The MEFs described in (**a**) were analyzed for proliferation over a 6-day time course in the presence or absence of 5 μM CX-4945.

### Supplementary Figure 5

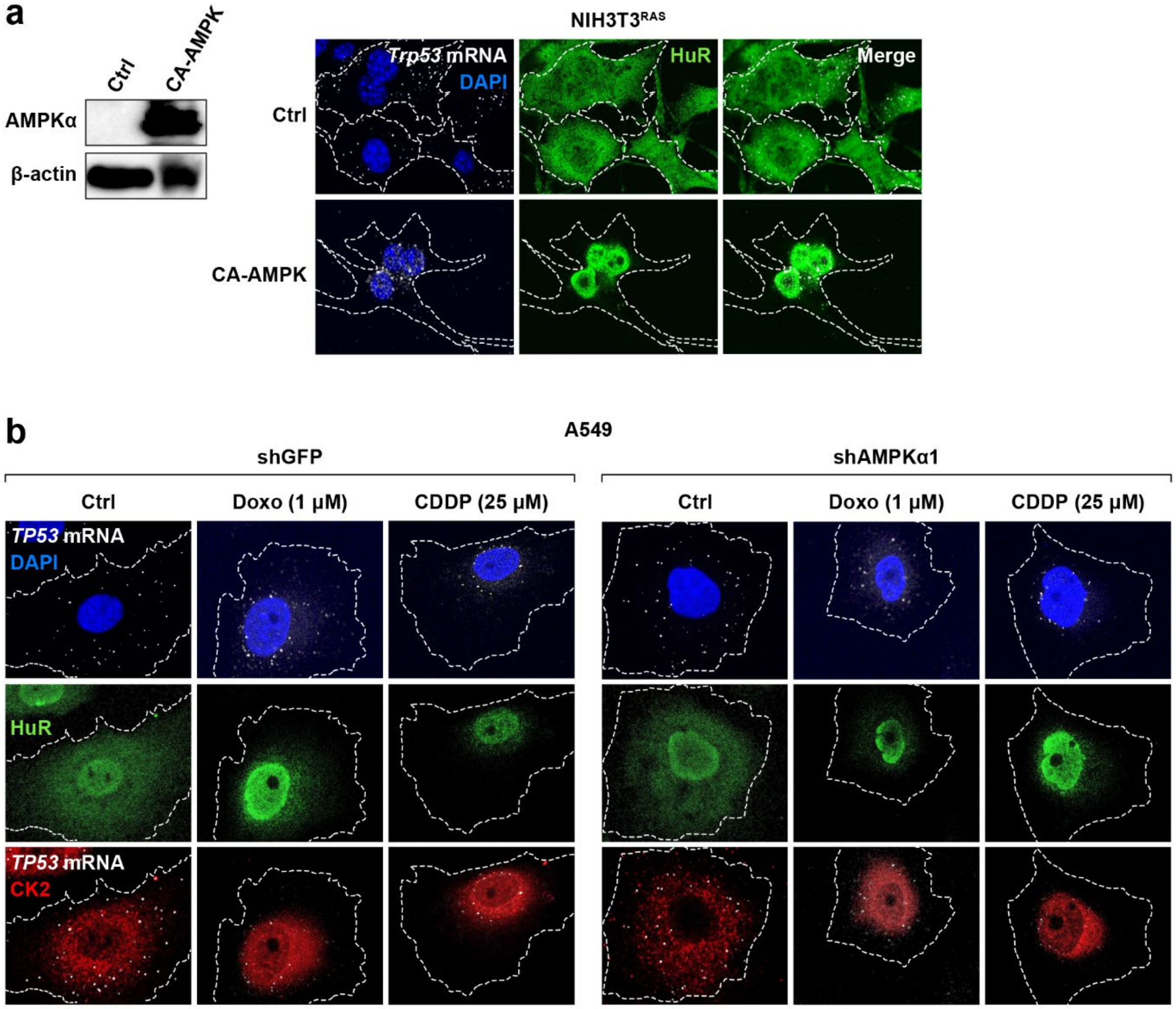


**Supplementary Fig. 5 Perinuclear translocation of *TP53* transcripts is induced by constitutively active AMPK but is independent of AMPKα1 in cells exposed to DNA damaging agents. a** CA-AMPK-expressing or control NIH3T3$^{RAS}$ cells were analyzed by RNA FISH for *Trp53* mRNAs and immuno-stained for HuR. **b** AMPKα1 depletion in A549 cells does not affect perinuclear localization of *TP53* transcripts in cells treated with doxorubicin or cisplatin. The experiment was performed as described in Fig. 7c, except that AMPKα1 was depleted.
